# Both genetic and social inheritance influence female aggressive and submissive behavior in a wild primate

**DOI:** 10.64898/2026.11.06.764731

**Authors:** Beniamino Tuliozi, Emily M. McLean, Elizabeth A. Archie, Jenny Tung, Susan C. Alberts

**Affiliations:** Department of Biology, Duke University, Durham, NC 27708, USA; Division of Natural Sciences and Mathematics, Oxford College, Emory University, Oxford, GA 30054, USA; Department of Biological Sciences, University of Notre Dame, Notre Dame, IN 46556, USA; Department of Primate Behavior and Evolution, Max Planck Institute for Evolutionary Anthropology, Leipzig 04103, Sachsen, Germany; Faculty of Life Sciences, Institute of Biology, Leipzig University, Leipzig 04103, Sachsen, Germany; Department of Evolutionary Anthropology, Duke University, Durham, NC 27708, USA

**Author notes:** These authors contributed equally to this work (Beniamino Tuliozi, Emily M. McLean). Corresponding author: Beniamino Tuliozi, Address: Duke University, Department of Biology, Box 90383, Durham, NC 27708, USA.

**Keywords:** Social behavior, genetic variance, social inheritance, aggressive behavior, indirect genetic effects, quantitative genetics

## Abstract

Agonistic interactions are important determinants of resource access in wild animals, but their genetic basis remains poorly understood. Here we use a quantitative genetics approach to estimate the heritability of aggressive and submissive behavior of wild female baboons, considering relative rates of both individual and dyadic aggression and submission. To control for the social inheritance that creates kin-based dominance hierarchies, we implemented a multi-matrix model that included a social status similarity matrix alongside the additive genetic matrix. Despite a substantial contribution of social inheritance, especially for submissive behavior, we found heritable components for both aggressive and submissive behavior (h^2^=0.21 and h^2^=0.09-0.11, respectively). In dyadic models, aggressive behavior was primarily influenced by the aggressor’s genotype (a direct genetic effect), while submissive behavior was primarily affected by the social partner’s genotype (an indirect genetic effect). Finally, we found evidence for a negative genetic (but not phenotypic) correlation between aggressive and submissive behavior within dyads, and a negative genetic correlation between submissive behavior and eliciting submissive behavior. Together, our results highlight how direct genetic, indirect genetic, and social inheritance effects combine to influence primate social behavior.

## INTRODUCTION

Agonistic interactions comprise the suite of aggressive and submissive behaviors that animals use to enact social conflict. These interactions are often species specific and have profound effects on individual fitness: agonisms are typically risky and energetically costly, but animals who win agonistic interactions usually benefit by gaining access to resources (Ellis 1995; Stockley & Bro Jørgensen 2011; Clutton Brock & Huchard 2013; Huchard & Lukas 2022). If repeatable and consistent within groups, the outcomes of dyadic interactions can also give rise to dominance hierarchies that structure resource access over a longer period (reviewed in Strauss *et al*. (2022). Because of their clear fitness relevance, ggressive behaviors have been the subject of intense interest for decades (e.g. Collias (1944); Tinbergen (1968); Holekamp and Strauss (2016). Submissive behaviors, on the other hand, have received much less attention but are as important as aggressive behaviors for determining the outcomes of social conflicts (reviewed in Reddon *et al*. 2021). Given the strong link between agonistic behavior and fitness, it is no surprise that much work has been devoted to understanding selection on, and the evolution of, agonistic behaviors (Smith & Harper 1988; Moore *et al*. 2002; Lindenfors & Tullberg 2011; Tobias *et al*. 2012; Holekamp & Strauss 2016).

However, animal populations can only respond to selection on agonistic behavior if it has a partially heritable (i.e., genetic) basis. Several lines of research have provided evidence of a heritable basis for agonistic behaviors in vertebrates. Research in commercial populations of livestock has shown that agonistic and aggressive behaviors have low, but non-zero heritability (Løvendahl *et al*. 2005; Turner *et al*. 2008; Sartori & Mantovani 2010; Stukenborg *et al*. 2012), with studies on Aosta cattle show that multi-generational selection on social dominance can cause evolutionary change in a population’s morphology and physiology (Tuliozi *et al*. 2022; Tuliozi *et al*. 2023). Research on captive or temporarily trapped wild animals also supports the heritability of agonistic behavior, typically in the context of mirror tests or experimentally staged agonistic encounters between pairs of individuals (e.g., Wilson *et al*. (2009); Taylor *et al*. (2012); Araya Ajoy and Dingemanse (2017); Murlanova *et al*. (2022). An advantage of experimental tests of this sort is that they provide a standardized metric of aggression in a relatively large number of animals. At the same time, however, they also exclude the ecological and social contexts in which agonistic behaviors are naturally expressed. This limitation makes the generalizability of such results to natural settings unclear (Biro 2012; Osborn & Briffa 2017; Kondrakiewicz *et al*. 2019).

Studies of the genetic basis of naturally occurring agonistic interactions in the wild are therefore crucial for understanding the evolution of social behavior. These studies remain scarce, but those that exist generally support the idea that some aspects of naturally occurring agonistic behaviors are heritable in natural or semi-natural settings. For instance, substantial heritability has been reported for aggression in a free-ranging population of rhesus macaques (Brent *et al*. (2013), while low-to-modest heritability has been reported for agonistic network traits in yellow-bellied marmots (Lea *et al*. (2010), aggressiveness in spotted hyenas (Yoshida *et al*. (2016), and social dominance in red deer (Wilson *et al*. (2011).

These findings suggest that quantitative genetic dissection of agonistic behavior is possible in wild populations of social animals. However, at least three gaps in our understanding remain. First, agonistic behavior typically involves dyadic interactions, raising the possibility that the genotype of a focal animal’s partner might also influence the focal’s tendency to exhibit either aggressive or submissive behavior (Wilson *et al*. 2009; Lea *et al*. 2010; Santostefano *et al*. 2017). This type of effect, known as an indirect genetic effect (IGE) or sometimes “social genetic” effect, is well-documented in captive animals (Saltz 2013; Baud *et al*. 2017), but few studies have considered the contribution of both IGEs and DGEs in natural populations (Fisher *et al*. 2019; Santostefano *et al*. 2021; McLean *et al*. 2023; Spitz *et al*. 2026). If IGEs are substantial, they can increase the genetic contribution to trait variance (sometimes well above that attributable to DGEs), and the magnitude and direction of correlations between DGEs and IGEs can modulate the efficacy of selection on a fitness-related trait (Wolf *et al*. 1998; Bijma & Wade 2008; McGlothlin *et al*. 2010; Wilson 2014; Girardeau *et al*. 2025).

Second, most quantitative genetic studies of agonistic interactions have focused on aggressive behaviors, or on social dominance interpreted as a win-lose dyadic trait (Wilson *et al*. 2011; Tuliozi *et al*. 2023). Despite their importance to dominance outcomes, previous studies in natural populations have not yet examined genetic effects on submissive behavior (either direct or indirect) or the genetic correlation between submissive and aggressive behavior. At the phenotypic level, a recent study on wild spotted hyenas (*Crocuta crocuta*) found that aggressive and submissive behaviors were not correlated, suggesting that they do not simply reflect two oppositely coded expressions of the same trait (McCormick & Holekamp 2022). In support of this possibility, the neural pathways involved in submissive and aggressive behavior show little overlap in laboratory rodents, with submissive behavior having more in common with anti-predator responses than with aggressive behavior (Adams 2006; Murlanova *et al*. 2022). A lack of a genetic correlation between aggressive and submissive behavior would suggest that the two traits can evolve independently, as variation in aggressive behavior and submissive behavior might have distinct fitness consequences (McCormick & Holekamp 2022; Smit 2024).

Third, while a dual model of genetic and social inheritance of behavior has long been advocated in theoretical studies (Boyd & Richerson 1988), very few applications of this model exist for non-human animals (Danchin *et al*. 2013; Thomson *et al*. 2018). And yet, in some animal societies, including many primates and spotted hyenas, dominance rank – as established through agonistic interactions – is socially transmitted from mother to daughter in the form of a nepotistic dominance hierarchy (Holekamp & Smale 1991; Chapais 1992; Silk 2009). In these societies, dominance rank (social status) may thus be transmitted with high fidelity over many generations, independently of genetic inheritance (Engh *et al*. 2009; Langergraber 2012; Lea *et al*. 2014). For instance, in some baboon populations, daughters inherit the rank below their mother and usually above their older sisters (Lea *et al*. 2014). An individual’s success in a fight thus reflects not only her own competitive ability but also the relative social rank of her family. It follows that an individual’s agonistic behavior is likely to be shaped by the social inheritance of her dominance rank, as well as by the genetic inheritance. Because social inheritance can inflate estimates of heritability, differentiating it from genetic inheritance is a key unresolved issue in studies of both humans and non-human animals (Sheppard *et al*. 2018; Young *et al*. 2019; Nichols *et al*. 2021; Tan *et al*. 2024).

Here, we investigated the genetic basis of naturally occurring agonistic behavior in a wild primate population, the baboons of the Amboseli basin in southern Kenya. We estimated the heritability of both aggressive and submissive behaviors using the “animal model” (a traditional approach to quantitative genetic modeling: (Kruuk 2004; Wilson *et al*. 2010). We addressed the problem of social inheritance in our study system by including not only an additive genetic distance matrix, but also a social distance matrix (i.e., social status matrix) which captures the variance associated with the “social status effect” of occupying close positions within the dominance hierarchy (Stopher *et al*. 2012; Thomson *et al*. 2018). The social status matrix allowed us to compare how much of the variance in the phenotype is explained by similarity in social status (i.e., similarity in dominance rank) with how much is explained by genetic relatedness. We then used bivariate analyses to test whether aggressive and submissive behaviors are genetically correlated. Finally, we examined indirect genetic effects on both aggressive and submissive behaviors.

## METHODS

### Study population

The baboon population in the Amboseli basin of southern Kenya has been the subject of ongoing research for over five decades (Alberts & Altmann 2012; Alberts 2019). The ancestry of baboons in this population is primarily yellow baboon (*Papio cynocephalus*), but all individuals in the population contain low to moderate levels of genetic admixture from a congener, *P. anubis* (Alberts & Altmann 2001; Vilgalys *et al*. 2022). All animals in a subset of social groups (“study groups”) are individually recognized on sight, and both behavioral and demographic data are collected as part of the near-daily monitoring of the study groups (Alberts *et al*. 2020).

Our study subjects were all adult female baboons present in study groups between January 1983 and June 2023. Adult females are those that have reached menarche, which is visually apparent to observers and occurs at a median age of 4.5 years in the study population (Onyango *et al*. 2013). The behavioral dataset includes 165,630 agonistic interactions collected during the lives of 365 adult females, with a median of 216 interactions per individual (range 1-4477 interactions).

The research in this study was approved by the Institutional Animal Care and Use Committee (IACUC) at Duke University (A273-17-12) and University of Notre Dame (25-02-9112), by the Ethics Council of the Max Planck Society (2022_29) and adhered to the laws and guidelines of the Kenyan governments.

### Behavioral data collection

Our behavioral data consisted of counts of decided agonistic interactions between adult female baboons. Decided agonistic interactions are dyadic interactions that occur when one animal (the “loser” of the interaction) gives only submissive gestures and the other one (the “winner” of the interaction) gives aggressive or neutral gestures, but not submissive ones (Supplementary Table S1; for more details, see Alberts *et al*. (2020) and Hausfater (1974)). Occasionally, we observe dyadic interactions that are “undecided,” i.e., interactions in which one or both participants gives a mix of aggressive and submissive gestures. Because these constitute a small fraction of recorded interactions, we excluded them from this analysis.

We classified agonistic interactions as either (1) “Aggressive-Submissive” (hereafter AS) interactions, i.e., those in which the loser gave only submissive gestures and the winner gave only aggressive gestures, and (2) “Neutral-Submissive” (hereafter NS) interactions, i.e., those in which the loser gave unprovoked submissive gestures, or in response to a non-aggressive gesture by the winner (e.g., gazing, approaching, reaching).

Observers recorded agonistic interactions during “representative interaction sampling,” in which they moved systematically through the group conducting 10-min-long focal animal samples in a randomized order, simultaneously recording all agonistic interactions in their line of sight (including those that did not involve the subject of focal sampling) (Alberts & Altmann 2012; Alberts *et al*. 2020; Campos *et al*. 2020). We consider the resulting counts of agonistic interactions to be unbiased measures of the relative frequencies with which different individuals engage in agonistic behavior. Differences in group size can affect the relative frequencies of agonistic interactions because the number of observers remains relatively constant while group size varies over time. To correct for this effect, we further adjusted all metrics of agonistic interactions by regressing them against daily rates of observer effort, defined as the number of focal animal samples collected in a group per adult female per day (Supplementary Methods S1, Campos *et al*. (2020); McLean *et al*. (2023).

### Agonism indices

We created two types of agonism indices (aggregate and dyadic), following an approach we previously used to investigate grooming behavior in this population (McLean *et al*. (2023); see below and Supplementary Methods S2-S3 for additional details). Each index was calculated for each subject, separately for aggressive and submissive behaviors, on a yearly basis, resulting in repeated measures for most subjects. Index values were then used as response variables in quantitative genetic animal models (see “Pedigree, animal models, and fixed effects” section).

#### Aggregate indices

The aggregate indices represent the relative frequency with which each female was observed to participate in decided agonistic interactions, compared to all other adult females in the population during the same year (Figure 1). The aggregate indices allowed us to investigate individual propensities for engaging in agonistic behaviors, irrespective of opponent identity. We calculated three aggregate indices for each female, in each year of her adult life starting from her birthday: 1) The index of *aggregate aggression given* by the focal female to all opponents, using AS interactions; 2) the index of *aggregate submission given to aggressive opponents* by the focal female, using AS interactions; and 3) the index of *aggregate submission given to neutral opponents* by the focal female, using NS interactions (Figure 1). A positive value for an index in a given year therefore reflects a female who was observed to perform agonistic gestures at a higher relative frequency than other females in the population in that year, controlling for observer effort. Negative values correspond to females who were observed to perform such gestures at a lower relative frequency in that year, controlling for observer effort. (for additional details see Supplementary Methods S2, Supplementary Figure S1).

**Figure 1.**
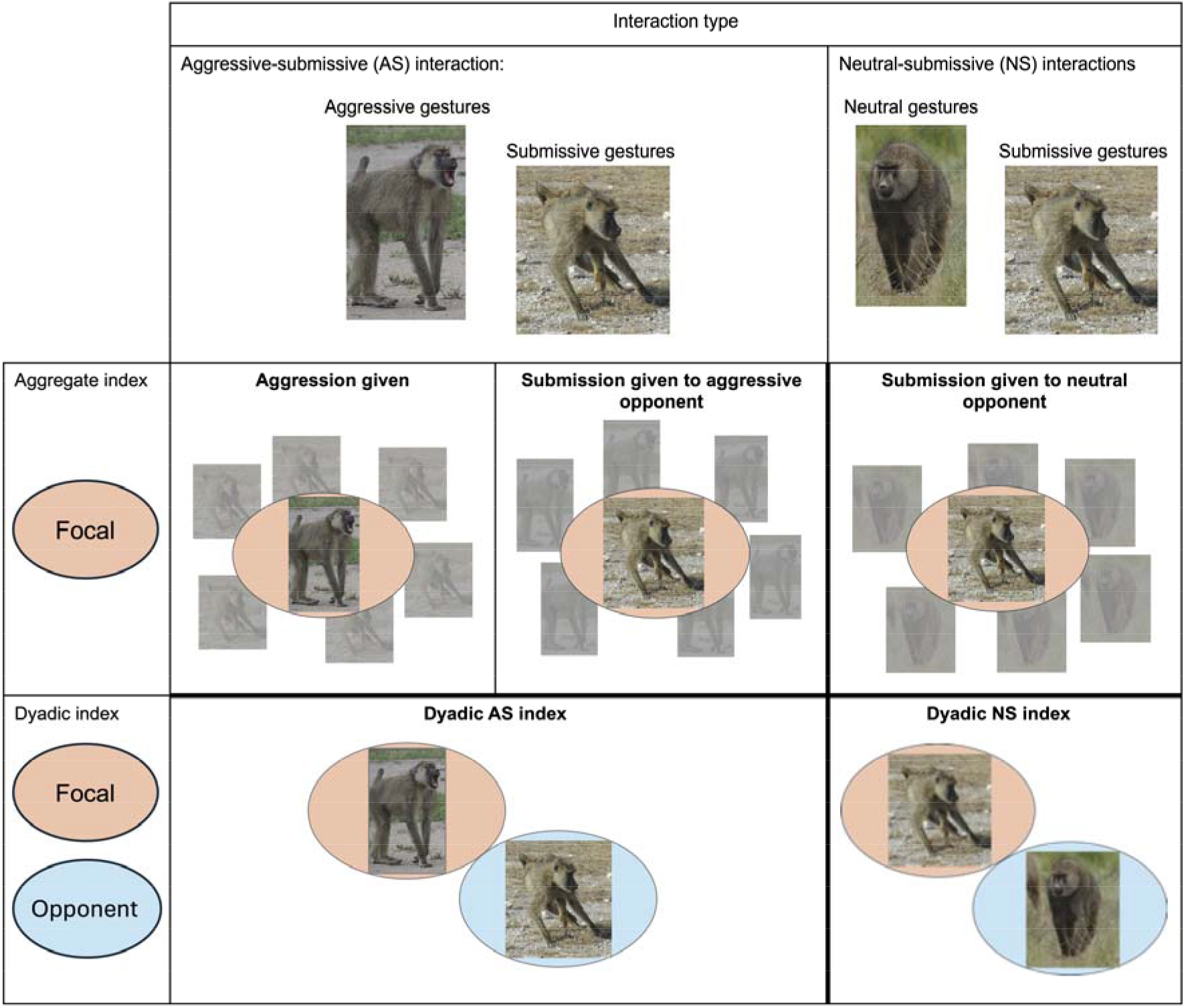
Graphical representation of agonism indices. Columns represent the agonistic interaction constituting the index (aggressive-submissive or neutral-submissive). Rows represent the type of index: aggregate indices (built based on all interactions involving a focal female in a given year of her life) and dyadic indices (built based on all interactions between two females in a given calendar year). Ovals represent the role of the individuals in the models: pink ovals represent focal individuals and light blue ovals represent opponents.

#### Dyadic indices

The dyadic indices were calculated for each pair of co-resident adult females in the population, i.e., females that lived in the same group at the same time for at least 60 days during the calendar year and had at least one decided agonistic interaction. These indices represent the frequency with which each dyad was observed to have a decided agonistic interaction in a given calendar year, relative to the frequency with which all other dyads were observed to have decided agonistic interactions in the same time interval, controlling for observer effort. The dyadic indices allowed us to investigate individual propensities to engage in agonistic interactions that (unlike the aggregate indices) were contingent on the identity of the opponent (Figure 1; see also Supplementary Methods S3).

Each dyadic interaction had a focal individual – the individual contributing the direct genetic component to the phenotypic variance – and an opponent – the individual contributing the indirect genetic component to the phenotypic variance. Thus, the dyadic indices investigate how the identity of both focal (direct genetic effect) and opponent (indirect genetic effect) influenced the behavior of the focal individual via genetic effects, whether the focal’s behavior was aggressive or submissive.

We calculated two dyadic indices, both of which are illustrated in Figure 1: 1) The *dyadic AS index* represents the relative frequency of interactions between a given pair in which the focal gave aggressive gestures and the opponent gave submissive gestures, within a given calendar year. 2) The *dyadic NS index* represents the relative frequency of interactions between a given pair of individuals in which the opponent gave no aggressive gestures but the focal gave submissive gestures, within a given calendar year. We did not pool AS and NS interactions to model submissive behavior because, although submissive gestures are performed in both cases, the contribution of the opponent is different between the two types of interaction (in one case, the opponent is aggressive, and in the other case the opponent is neutral), making a combined index more difficult to interpret.

For each pair of individuals that interacted in each year we calculated separate AS and NS dyadic indices for each direction. That is, we calculated one AS index based on interactions where individual 1 was the focal (i.e, gave aggressive gestures) and individual 2 was the opponent (i.e., gave submissive gestures), and a separate AS index based on interactions where individual 2 was the focal (gave aggressive gestures) and individual 1 was the opponent (gave submissive gestures). We followed the same procedure for the NS index. Positive values of the dyadic index indicate cases in which a given focal/opponent dyad had high frequencies of “AS” or “NS” interactions, relative to all other dyads in the population for that year, while negative values indicate cases in which a given dyad had relatively low frequencies of that interaction type (for additional details see Supplementary Methods S3, Supplementary Figure S1).

### Pedigree, animal models, and fixed effects

We combined pedigree information with phenotypic and environmental data in a series of mixed effects models (“animal models”) that allowed us to partition the phenotypic variance of all our measures of agonistic behavior into additive genetic and other variance components (Lynch & Walsh 1998; Kruuk 2004). All our models were fitted using the ‘asremlr’ package (Gilmour *et al*. 2009) in the R environment (R Core Team 2013). All results presented below are based on models that converged and where visual checks of model residuals revealed no obvious violations of model assumptions.

Estimating genetic effects requires information about relatedness within the sample, which we generated using a multigenerational pedigree based on birth records and genetic parentage assignment carried out using 7-14 microsatellite genotypes. The maternities of all our study subjects were known, but only 66% of the paternities, which require genetic material from both offspring and potential fathers, including the true father, were known. Paternity was based on genetic exclusion and further supported through the use of the likelihood-based paternity assignment program CERVUS 2.0, with confidence set at 95% (see Buchan *et al*. (2003); Alberts *et al*. (2006); Van Horn *et al*. (2007) for methodological details). The full pedigree includes more than 1,500 individuals (Galezo *et al*. 2022) but the subset of this pedigree necessary to describe the relationships between all 365 of our study subjects consists of 473 individuals (for pedigree details see Supplementary Table S2).

In each of our animal models we included a series of fixed effects representing non-genetic factors known or predicted to influence agonistic behavior (see Supplementary Table S3 for details). For the aggregate agonism indices, we included fixed effects for: (i) average dominance rank of the focal during the year in which the index was measured, (ii) age of the focal in that year, (iii) average size of the focal’s social group during that year, and (iv) average adult sex ratio in the focal’s social group during that year. To test if the presence and number of relatives in the group influenced agonistic behavior, we also fitted fixed effects of the proportion of time in each year the focal lived in the same group as her (v) mother and (vi) father (if known), and the sum of the proportions of times in the year that the focal co-resided with, respectively, one of her (vii) daughters, (viii) sisters, (ix) or adult sons. The three latter metrics could range from 0 as a minimum (no co-resident relatives in the observation year) to a maximum of N, where N is the total number of the focal’s daughters, sisters (half or full-sib), or adult sons known to us. A value of 0 would mean that none of those relatives co-resided with the focal during the year, while a value of N would mean that all known relatives in these kin classes co-resided with the focal female for the full year. Partial year co-residency was taken into account as a fractional value of the year.

In the dyadic models, we again included fixed effects for dominance rank, age, group size, and sex ratio (see items i - iv above). We also included: (v) dominance rank of the opponent, (vi) interaction between the focal’s and the opponent’s dominance ranks, (vii) age of the opponent, (viii) interaction between the focal’s and the opponent’s ages, (ix) whether the dyad was a mother-daughter pair or (x) a maternal sister pair, and (xi) the pedigree relatedness between focal and opponent (Supplementary Table S4).

To account for the possibility that genetic admixture between *P. cynocephalus* and *P. anubis* influences agonistic behavior, we also ran models that included an estimate of the individual’s degree of genetic admixture as a fixed effect (Vilgalys *et al*. 2022). Because we only have genetic ancestry scores for 45% of the individuals in this dataset, we report this analysis only in the supplement. Briefly, the variance estimates recovered from the models including genetic ancestry were qualitatively similar to those reported in the main results (Supplementary Methods S4; Supplementary Table S5).

### Accounting for the social environment through an S matrix approach

Dominance rank in female baboons in Amboseli is socially inherited with high fidelity, such that daughters typically insert into the linear adult female dominance hierarchy below their mothers and above their older sisters (Chapais 2002, Silk 2009). In this population, this process results in long periods of inter-generational stability in the relative ranks of female matrilines, although the general pattern of stability can sometimes be punctuated by episodes of change (Hausfater *et al*. 1982; Samuels *et al*. 1987; Lea *et al*. 2014). Failing to account for this strong, non-genetic source of familial resemblance is likely to result in inflated estimates of quantitative genetic parameters such as heritability. To capture the variance in agonistic behavior associated with social inheritance of dominance rank, we therefore implemented a multi-matrix form of the animal model (Stopher *et al*. 2012; Thomson *et al*. 2018) that is often used to account for spatial similarity (Rutschmann *et al*. 2020; Jones *et al*. 2024; Satarkar *et al*. 2025). The classic animal model uses an additive genetic relatedness matrix (**A** matrix) to calculate how much phenotypic variance is explained by genetic similarities between two individuals. Our multi-matrix animal model includes an **S** (“social status”) matrix in addition to the **A** matrix, which accounts for social status similarity between individuals (Stopher *et al*. 2012; Thomson *et al*. 2018).

Specifically, we constructed the social status **S** matrix by describing the degree of similarity in dominance rank between every pair of females in our study. The **S** matrix reflects pairwise differences in lifetime average dominance rank, capturing the percentage of variance associated with the “social status effect” of occupying close positions within the dominance hierarchy. The **S** matrix therefore quantifies how much of the variance in the phenotype can be attributed to variance in this lifetime measure of social status. The resulting vector of estimated “status values” has a variance-covariance structure determined by the **S** matrix. These status values are analogous to breeding values, in that they represent the shift from the population phenotypic mean of an individual that can be attributed to its social status. In other words, variance in the random effect of social status captures the variance associated with similarity in dominance rank. For conceptually similar approaches see also Frère *et al*. (2010); Danchin *et al*. (2013); Nichols *et al*. (2021).

We calculated the distance in social status between individuals *i* and *j* as *e*^-|*si*-*sj*|^ where is the ordinal dominance rank of female *i* averaged over all her adulthood and *s_j_* is the ordinal dominance rank of female *i* averaged over the entire adulthood of individual *j* (Bhattacharyya 1943; Stopher *et al*. 2012; Thomson *et al*. 2018). Female ordinal rankings are calculated monthly by building a symmetrical NxN matrix containing all N females, where each cell contains the number of decided agonisms where the individual in the row won against the individual in the column. Ordinal rank is calculated by ordering rows to minimize the number of entries appearing below the diagonal of the matrix (Gordon et al., 2023). Importantly, female dominance rank is quite stable over the lifetime: less than 50% of adult females ever actively change their relative rank, and such changes are typically just one or two ordinal positions. The vast majority of the changes in ordinal dominance rank are instead due to females dying (exiting the hierarchy) or maturing (entering the hierarchy).

Although both the fixed effect of dominance rank and the **S** matrix are based on measures of ordinal dominance rank, we note that we use the term “dominance rank” to refer to the linear fixed effect of ordinal dominance rank below, and the term “social status” to refer the random effect associated with the position in the dominance hierarchy. This terminology is intended to differentiate the linear effect of rank on the phenotype from behavioral similarities (i.e., covariance) arising from similar social status and related attributes (e.g., in some species, individuals with similar ranks can also share more similar social networks: (Ilany *et al*. 2021).

### Analysis of aggregate agonism indices

For each of our three aggregate agonism indices (aggression given, submission given to an aggressive opponent, and submission given to a neutral opponent) we investigated genetic and non-genetic sources of variance in the aggregate behavior of the focal individual by constructing a series of linear mixed models with consistent fixed effect structures and random effect structures of serially increasing complexity (see the full series in Supplementary Methods S5, equations S1-S4). The aggregate indices were treated as approximately normal based on visual inspection of their distributions (Supplementary Figure S1) and of the distributions of model residuals. We used a likelihood ratio test to determine the best-fit model for each aggregate index. For all three aggregate indices, the best model included both an additive genetic component, indicating non-zero heritability, and a random effect of social status based on the **S** matrix). We refer to this formulation as the “heritability & status value model:”

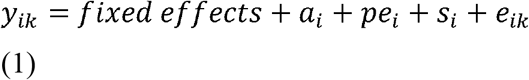

where *y_ik_* is the aggregate agonism index for (focal) individual *i* in year *k*, *e_ik_* is a residual error term, *a_i_* is the additive genetic contribution of individual *i* (i.e., her breeding value), *pe_i_* represents the permanent environment effect of individual *i*, and *s_i_* is the social status contribution of individual *i* (i.e., the **S** matrix analogue of the breeding value). For all models, we obtained estimates of the variance attributed to each component. We used the estimates for the additive genetic variance (σ^2^*_a_*) to calculate narrow-sense heritability (h^2^ = σ^2^*_a_* / σ^2^*_P_*), and the estimates for variance associated with social status (σ^2^*_S_*) to calculate the “social heritability” value (s^2^ = σ^2^*_S_* / σ^2^*_P_*), where σ^2^*_P_* is the total phenotypic variance.

We also constructed bivariate animal models to investigate the covariances among the random effects (genetic, permanent environment, and social) estimated for 1) aggregate aggression given and aggregate submission given in response to an aggressive opponent and 2) aggregate submission given to an aggressive opponent and aggregate submission given to a neutral opponent. To estimate each covariance, we constructed separate bivariate models, each with a different unstructured variance-covariance matrix (for breeding value and/or status value and/or permanent environment effects). To test the significance of each potential source of covariance (genetic, social, permanent environmental), we used a likelihood ratio test to compare each of these models to a model in which all covariances were constrained to zero and only the residual covariance between traits was allowed to vary. For the full variance-covariance matrix structure of our bivariate models see Supplementary Methods S6. For both bivariate models we also estimated the phenotypic correlations between pair of aggregate indices. We also compared the phenotypic correlations estimated in these models to an alternative specification, where we fitted bivariate models including only the effect of individual identity without further partitioning between genetic, social and permanent environmental effects, and calculated both the individual-level correlation and the phenotypic correlation between the traits.

### Analysis of dyadic agonism indices

For each of our dyadic agonism indices (the dyadic AS index and the dyadic NS index) we constructed a series of linear mixed models with consistent fixed effects structures and increasingly complex random effect structures, which included both the additive genetic variance matrix and the social status matrix (the full series is presented in Supplementary Methods S5, equations 5-10). The dyadic indices departed from normality, showing a right-skewed distribution (Supplementary Figure S1). We chose to fit a linear model to these indices, as such models are relatively robust to departures from normality (Schielzeth *et al*. 2020; Knief & Forstmeier 2021), but compared the results estimated from our best-fitting dyadic models (see below) with the results estimated from the same models run on log-transformed dyadic indices.

Unlike the aggregate indices, the dyadic index allowed us to estimate both direct and indirect genetic effects. Direct genetic effects (DGE) quantify the additive genetic effect of the focal individual’s genotype on its phenotype, while indirect genetic effects (IGE) quantify the additive genetic effect of the social partner (opponent’s) genotype on the focal individual’s phenotype (Wolf *et al*. 1998).

The best supported model for both the dyadic AS index and the dyadic NS index was the “Dyadic heritability model (DGE + IGE)”, which included both direct and indirect genetic effects, but not the **S** matrix (see Supplementary Methods S5, S7):

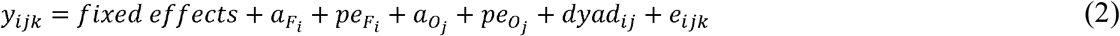

In both models, *y_ijk_* is the frequency of interactions between the focal *i* and the opponent *j* in year *k.* In the AS index, the focal individual *i* gave aggressive gestures, while in the NS index the focal individual *i* gave submissive gestures. We also included a term, *dyad_ij_* (an identity assigned to each unique pair of individuals) to take into account repeated measures for each dyad. Here, *e_ijk_* is residual error, *a_F_i__* is the additive genetic contribution of the focal *i*, *pe_F_i__* is the permanent environment effect of the focal. In addition, 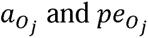 are respectively the additive genetic contribution and the permanent environmental effect of the opponent. In the AS index, the opponent is defined as a partner who performed a submissive gesture in response to aggressive gestures; in the NS index, the opponent is defined as a partner who performed no or neutral gestures in the interaction but was the receiver of submissive gestures.

Because direct and indirect genetic effects 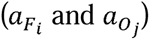 were left free to covary, we were able to estimate the direct genetic variance 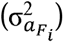, the indirect genetic variance 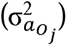, and the covariance between them 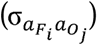. We rescaled the covariance to a correlation and used a likelihood ratio test with one degree of freedom to test if the correlation was significant by comparing the model in which the correlation was free to vary to models in which the correlation was constrained to 0 or 1. If the correlation was not significant, we excluded it from the model. We did the same for the permanent environmental covariance 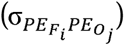 between focal permanent environment 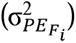 and opponent permanent environment 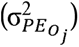 (Supplementary Methods S5, equations 11-12).

## RESULTS

### Aggregate indices: heritability and social status variance

The best model for each of the aggregate indices included random effects for both additive genetic variance and social status variance (‘Heritability & status value’ model in Table 1). Aggregate aggression given was moderately heritable (h^2^=0.21±0.05; Figure 2) while both of our submission indices were somewhat less so (submission given to aggressive opponent: h^2^=0.11±0.04; submission given to neutral opponent: h^2^=0.09±0.04; Figure 2).

**Figure 2.**
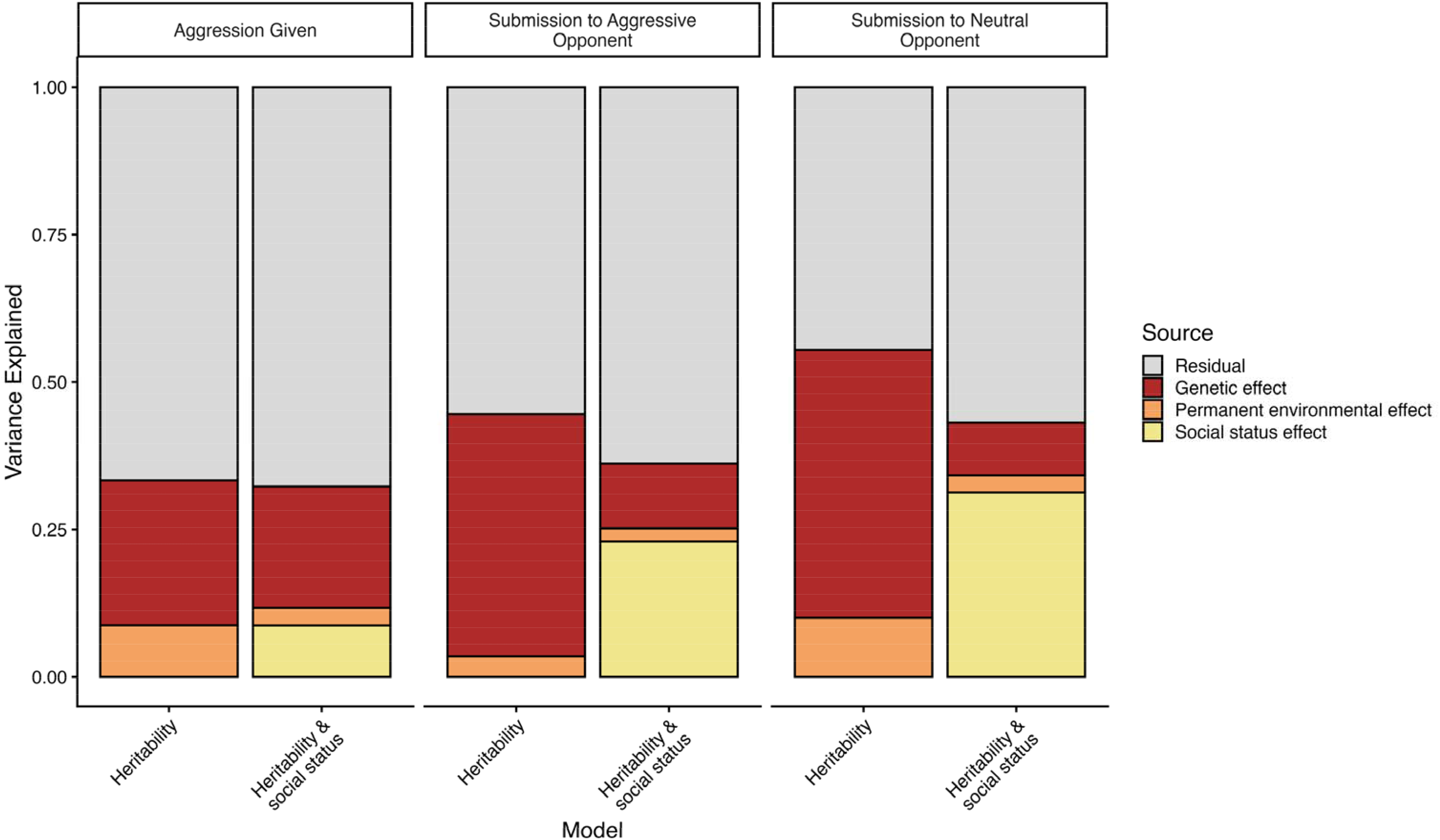
Proportions of variance explained by random effects in three aggregate indices of agonism. The two different models for each index (‘Heritability’, ‘Heritability and social status’) differ in whether the **S** matrix is included to account for the variance attributable to social inheritance.

**Table 1.** Aggregate models: Model comparisons for aggregate agonism indices, showing estimates of variance components with standard errors in parentheses below each estimate. Log-likelihoods (LnL) are shown for each model: all models represent a significantly (p < 0.001) better fit than the model above, with models in **bold** having highest log-likelihood for each index.

| Index | Model name<sup>a</sup> | σ<sub>a</sub><sup>2</sup> | σ<sub>PE</sub><sup>2</sup> | σ<sub>S</sub><sup>2</sup> | σ<sub>F</sub><sup>2</sup> | σ<sub>e</sub><sup>2</sup> | *h*<sup>2</sup> | *s*<sup>2</sup> | LnL |
|---|---|---|---|---|---|---|---|---|---|
| Aggression Given | Null model | -- | -- | -- | -- | 0.300 (0.008) | -- | -- | 233.11 |
|  | Repeatability model | -- | -- | -- | 0.101 (0.011) | 0.210 (0.006) | -- | -- | 469.25 |
|  | Heritability model | 0.077 (0.019) | 0.027 (0.013) | -- | -- | 0.209 (0.006) | 0.246 (0.054) | -- | 486.16 |
|  | **Heritability & status value model** | **0.064 (0.016)** | **0.009 (0.011)** | **0.027 (0.009)** | **--** | **0.209 (0.006)** | **0.206 (0.048)** | **0.087 (0.028)** | **497.52** |
| Submission Given to aggressive opponent | Null model | -- | -- | -- | -- | 0.441 (0.012) | -- | -- | -271.97 |
|  | Repeatability model | -- | -- | -- | 0.192 (0.019) | 0.262 (0.008) | -- | -- | 124.95 |
|  | Heritability model | 0.193 (0.036) | 0.016 (0.020) | -- | -- | 0.261 (0.008) | 0.411 (0.060) | --- | 151.61 |
|  | **Heritability & status value model** | **0.045 (0.017)** | **0.009 (0.014)** | **0.094 (0.019)** | **--** | **0.263 (0.008)** | **0.110 (0.040)** | **0.229 (0.038)** | **196.95** |
| Submission Given to neutral opponent | Null model |  |  |  |  | 0.449 (0.012) |  |  | -296.36 |
|  | Repeatability model |  |  |  | 0.255 (0.023) | 0.214 (0.006) |  |  | 324.35 |
|  | Heritability model | 0.217 (0.044) | 0.048 (0.026) | -- | --- | 0.213 (0.006) | 0.454 (0.071) |  | 350.95 |
|  | **Heritability & status value model** | **0.034 (0.014)** | **0.011 (0.012)** | **0.119 (0.021)** | **---** | **0.218 (0.006)** | **0.090 (0.035)** | **0.312 (0.042)** | **420.13** |

<sup>a</sup>See Supplementary Methods S5 for enumeration and specification of all nested models
σ<sub>a</sub><sup>2</sup>= additive genetic variance
σ<sub>PE</sub><sup>2</sup>= variance attributable to the permanent environment of the focal
σ<sub>S</sub><sup>2</sup>= variance attributable to the social status of the focal
σ<sub>F</sub><sup>2</sup> = variance attributable to the focal individual, applicable only to models that do not include additive genetic effects
σ<sub>e</sub><sup>2</sup> = residual variance
*h*<sup>2</sup>= proportion of variance explained by additive genetic variance (σ<sub>a</sub><sup>2</sup>)
*s*<sup>2</sup>= proportion of variance explained by social status variance (σ<sub>S</sub><sup>2</sup>)

The social status matrix (**S** matrix) played an important role in these models: including the **S** matrix resulted in a dramatic reduction in the heritability estimates for both of the aggregate submission indices, relative to models that did not account for social inheritance (Table 1). Specifically, the heritability (h^2^) of submission given to an aggressive opponent was 0.41 when social inheritance was not considered, but it dropped to 0.11 in the best-fitting model reported above, which included the **S** matrix (Figure 2). Similarly, the heritability of submission given to a neutral opponent was 0.45 in a model excluding the **S** matrix, but only 0.09 when the **S** matrix was included (Table 1; Figure 2). In contrast, although we also detected a non-zero social inheritance effect in our model of aggregate aggression given, adding the **S** matrix did relatively little to alter the heritability estimates for this trait (h^2^=0.25 without the **S** matrix, h^2^=0.21 with the **S** matrix, Table 1; Figure 2). Accordingly, the proportion of phenotypic variance attributed to social status variance (s^2^, the “social heritability” analogue of genetic heritability), as captured by the **S** matrix, was much lower for aggregate aggression given (s^2^=0.09) than for both aggregate submission given to an aggressive opponent (s^2^=0.23) and aggregate submission given to a neutral opponent (s^2^=0.31); Table 1, Figure 2.

### Aggregate indices: correlations between indices

Our bivariate models converged only when at least two of the three non-residual covariances (genetic covariance, permanent environmental and social covariance) were set to 0. Thus, all the following results were obtained with models where only the covariance in question was allowed to vary (alongside the residual covariance). We found no significant genetic, permanent environmental, or social correlation between aggregate aggression given and aggregate submission given to an aggressive opponent (all p-values > 0.1; Supplementary Table S6). In contrast, aggregate submission given to aggressive opponents versus to neutral opponents showed strong, positive genetic correlations and status value correlations (both r > 0.96, p<0.0001, Supplementary Table S6). Phenotypic correlations between aggregate aggression given and aggregate submission given were weak and positive (0.07 < r_p_ < 0.11; Supplementary Table S6), while phenotypic correlations between aggregate submission given to aggressive and neutral individuals were strong and positive (0.65 < r_p_ < 0.70: Supplementary Table S6).

### Dyadic indices: direct and indirect genetic effects on agonistic behavior

The best model for each of our dyadic agonism indices included both direct and indirect genetic effects (Table 2). These results indicate measurable indirect genetic effects of opponent identity on the amount of aggression (AS index) and submission (NS index) that a focal female gave within a dyad. Direct genetic effects explained 10.4% of the variance (direct h^2^ = 0.10; Table 2, Figure 3) in the frequency of AS interactions that occurred in a dyad (relative to other female-female dyads in the population), while indirect genetic effects explained only 2.4% (indirect h^2^ = 0.02). In contrast, direct genetic effects explained approximately 4.4% of the variance in how many NS interactions occurred in a dyad (where the focal was submissive) and indirect genetic effects explained 6.5% of the variance (Table 2, Figure 3). Adding the social status matrix did not apparently improve the model for either of our dyadic indices (Supplementary Results S7).

**Figure 3.**
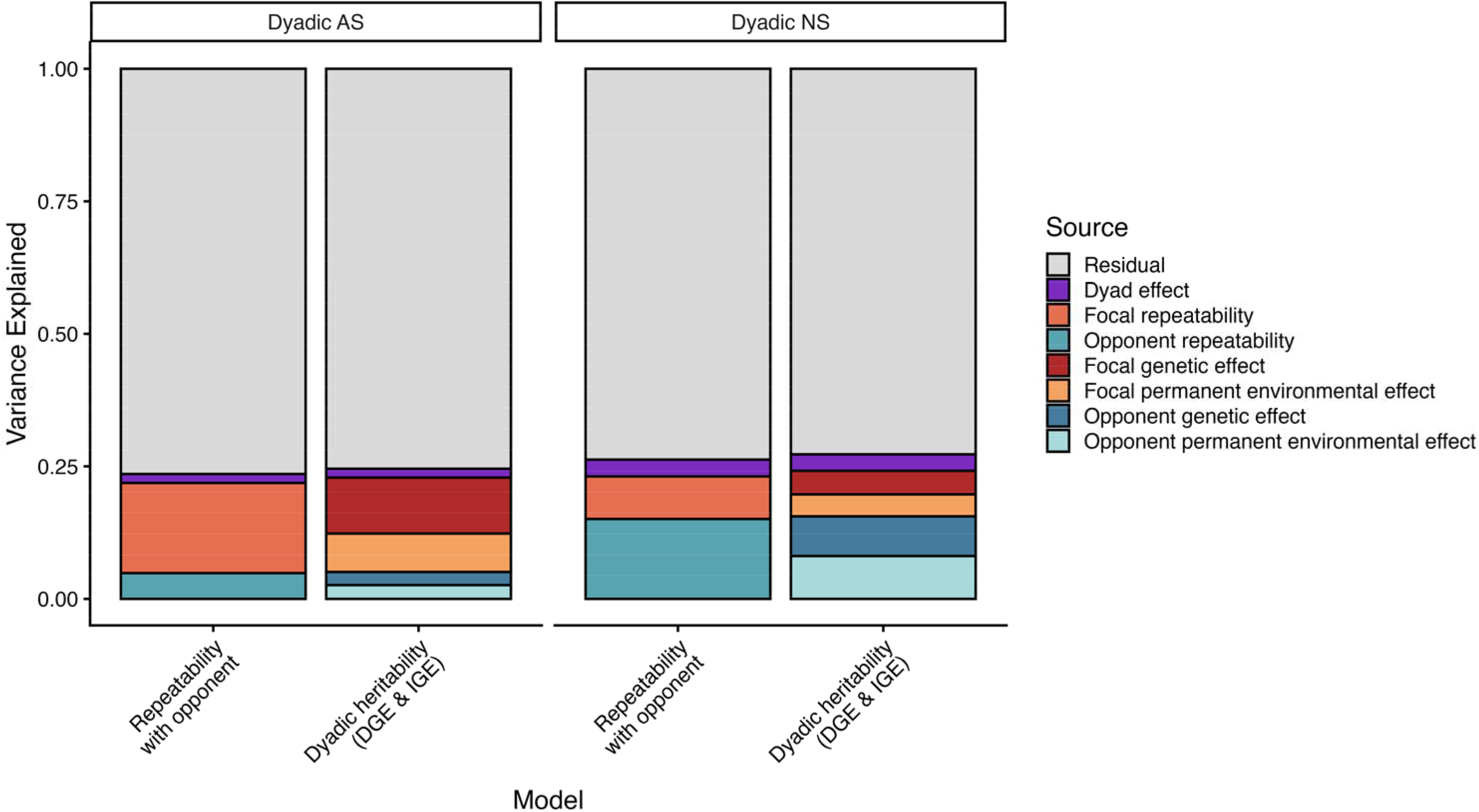
Proportions of variance explained by random effects in three dyadic indices of agonism. The two models shown for each index ‘Repeatability with opponent’ and ‘Dyadic heritability (DGE & IGE)’ differ in that the latter partitions individual variance (both of the focal and of the opponent) into additive genetic and permanent environmental variance.

**Table 2.**
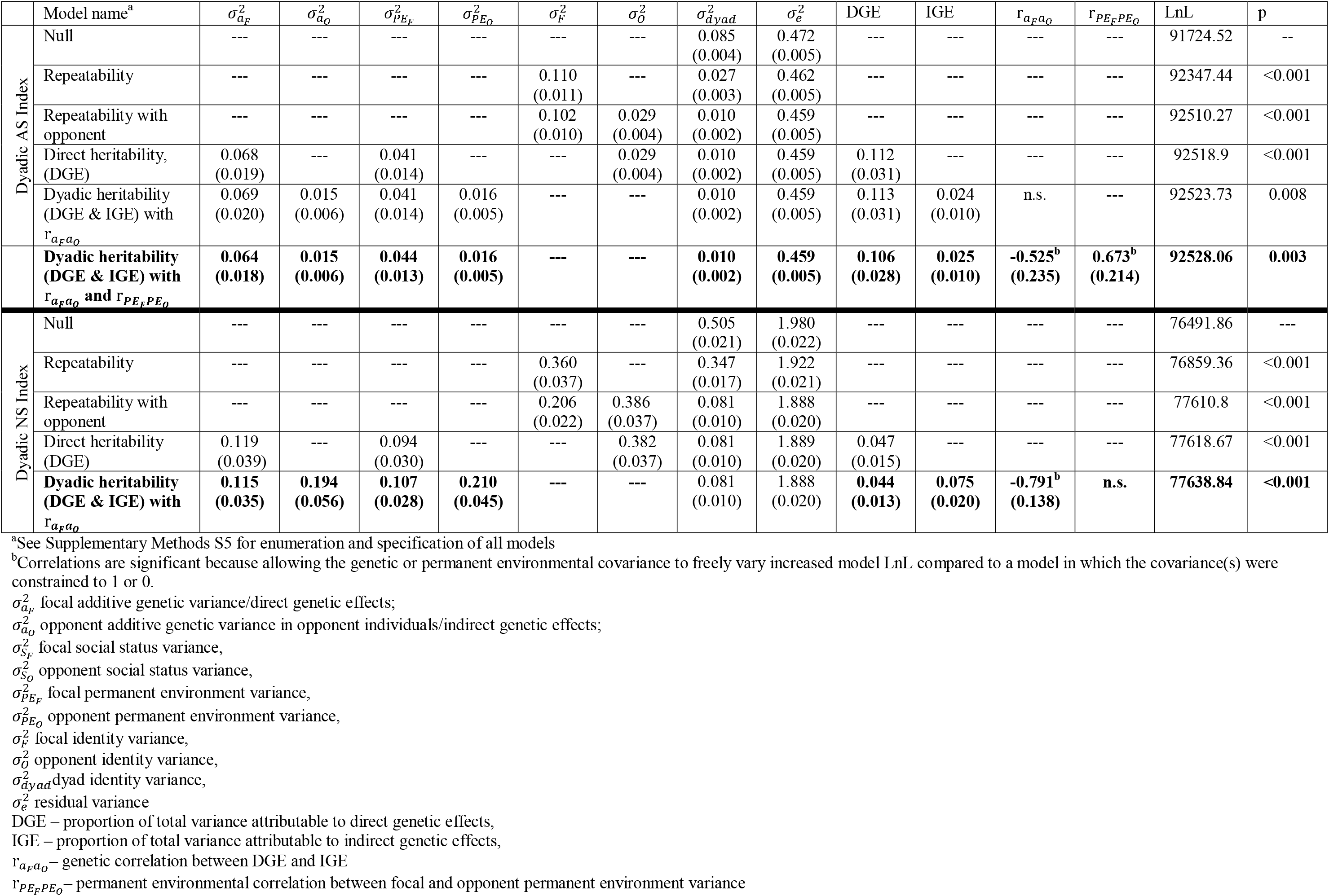
Dyadic models: Model comparisons for dyadic agonism indices, showing estimates of variance components (multiplied by a factor of 10^4^ to help visualization), with standard errors in parentheses below each estimate and log-likelihoods (LnL) for each model. P-values were determined by likelihood ratio tests, with degrees of freedom equal to the number of additional parameters under the more complex model, with models in **bold** having highest log-likelihood for each index.

|  | Model name<sup>a</sup> | $\sigma_{a_F}^2$ | $\sigma_{a_O}^2$ | $\sigma_{PE_F}^2$ | $\sigma_{PE_O}^2$ | $\sigma_{F}^2$ | $\sigma_{O}^2$ | $\sigma_{dyad}^2$ | $\sigma_{e}^2$ | DGE | IGE | r$_{a_Fa_O}$ | r$_{PE_FPE_O}$ | LnL | p |
|---|---|---|---|---|---|---|---|---|---|---|---|---|---|---|---|
| Dyadic AS Index | Null | --- | --- | --- | --- | --- | --- | 0.085<br>(0.004) | 0.472<br>(0.005) | --- | --- | --- | --- | 91724.52 | -- |
|  | Repeatability | --- | --- | --- | --- | 0.110<br>(0.011) | --- | 0.027<br>(0.003) | 0.462<br>(0.005) | --- | --- | --- | --- | 92347.44 | <0.001 |
|  | Repeatability with opponent | --- | --- | --- | --- | 0.102<br>(0.010) | 0.029<br>(0.004) | 0.010<br>(0.002) | 0.459<br>(0.005) | --- | --- | --- | --- | 92510.27 | <0.001 |
|  | Direct heritability, (DGE) | 0.068<br>(0.019) | --- | 0.041<br>(0.014) | --- | --- | 0.029<br>(0.004) | 0.010<br>(0.002) | 0.459<br>(0.005) | 0.112<br>(0.031) | --- | --- | --- | 92518.9 | <0.001 |
|  | Dyadic heritability (DGE & IGE) with r$_{a_Fa_O}$ | 0.069<br>(0.020) | 0.015<br>(0.006) | 0.041<br>(0.014) | 0.016<br>(0.005) | --- | --- | 0.010<br>(0.002) | 0.459<br>(0.005) | 0.113<br>(0.031) | 0.024<br>(0.010) | n.s. | --- | 92523.73 | 0.008 |
|  | **Dyadic heritability (DGE & IGE) with r$_{a_Fa_O}$ and r$_{PE_FPE_O}$** | **0.064<br>(0.018)** | **0.015<br>(0.006)** | **0.044<br>(0.013)** | **0.016<br>(0.005)** | **---** | **---** | **0.010<br>(0.002)** | **0.459<br>(0.005)** | **0.106<br>(0.028)** | **0.025<br>(0.010)** | **-0.525<sup>b</sup><br>(0.235)** | **0.673<sup>b</sup><br>(0.214)** | **92528.06** | **0.003** |
| Dyadic NS Index | Null | --- | --- | --- | --- | --- | --- | 0.505<br>(0.021) | 1.980<br>(0.022) | --- | --- | --- | --- | 76491.86 | --- |
|  | Repeatability | --- | --- | --- | --- | 0.360<br>(0.037) | --- | 0.347<br>(0.017) | 1.922<br>(0.021) | --- | --- | --- | --- | 76859.36 | <0.001 |
|  | Repeatability with opponent | --- | --- | --- | --- | 0.206<br>(0.022) | 0.386<br>(0.037) | 0.081<br>(0.010) | 1.888<br>(0.020) | --- | --- | --- | --- | 77610.8 | <0.001 |
|  | Direct heritability (DGE) | 0.119<br>(0.039) | --- | 0.094<br>(0.030) | --- | --- | 0.382<br>(0.037) | 0.081<br>(0.010) | 1.889<br>(0.020) | 0.047<br>(0.015) | --- | --- | --- | 77618.67 | <0.001 |
|  | **Dyadic heritability (DGE & IGE) with r$_{a_Fa_O}$** | **0.115<br>(0.035)** | **0.194<br>(0.056)** | **0.107<br>(0.028)** | **0.210<br>(0.045)** | **---** | **---** | 0.081<br>(0.010) | 1.888<br>(0.020) | **0.044<br>(0.013)** | **0.075<br>(0.020)** | **-0.791<sup>b</sup><br>(0.138)** | **n.s.** | **77638.84** | **<0.001** |

<sup>a</sup>See Supplementary Methods S5 for enumeration and specification of all models
<sup>b</sup>Correlations are significant because allowing the genetic or permanent environmental covariance to freely vary increased model LnL compared to a model in which the covariance(s) were constrained to 1 or 0.
$\sigma_{a_F}^2$ focal additive genetic variance/direct genetic effects;
$\sigma_{a_O}^2$ opponent additive genetic variance in opponent individuals/indirect genetic effects;
$\sigma_{S_F}^2$ focal social status variance,
$\sigma_{S_O}^2$ opponent social status variance,
$\sigma_{PE_F}^2$ focal permanent environment variance,
$\sigma_{PE_O}^2$ opponent permanent environment variance,
$\sigma_{F}^2$ focal identity variance,
$\sigma_{O}^2$ opponent identity variance,
$\sigma_{dyad}^2$dyad identity variance,
$\sigma_{e}^2$ residual variance
DGE – proportion of total variance attributable to direct genetic effects,
IGE – proportion of total variance attributable to indirect genetic effects,
r$_{a_Fa_O}$– genetic correlation between DGE and IGE
r$_{PE_FPE_O}$– permanent environmental correlation between focal and opponent permanent environment variance

The best-fitting AS index dyadic model included both genetic and permanent environmental covariances. We found a negative genetic correlation between the direct and indirect genetic effects of the AS index, suggesting that performing aggressive gestures is genetically negatively correlated with being the target of aggressive gestures (r*_a_F_a_O__* = -0.52±0.23, p=0.036; Table 2). The correlation between permanent environmental effects of focal and opponent was instead positive (r*_PE_F_PE_O__* = 0.67±0.21, p=0.003, Table 2).

We also found a strong, negative genetic correlation between the direct and indirect genetic effects on the NS index, indicating that performing unsolicited submissive signals is negatively genetically correlated with receiving unsolicited submissive signals (r*_a_F_a_O__* = -0.79±0.02, p<0.001; Table 2). In other words, individuals who tend to give submissive signals to other females, even in the absence of an aggressive interaction, also *receive* few unsolicited submissive behaviors from other females, and this correlation arises in part from a shared genetic basis. In the dyadic index, leaving the permanent environmental covariance free to vary did not improve the model (i.e., the permanent environmental correlation was not significant; r*_PE_F_PE_O__* = 0.14±0.21, p=0.498).

Using log-transformed dyadic indices did not result in qualitatively different variances; only the correlations moderately decreased in magnitude (Supplementary Table S7).

### Fixed effects

To facilitate comparison between the influence of predictor variables on each of the aggregate indices, as well as between the dyadic and aggregate indices, we report the magnitude of the fixed effects both in terms of percentage points (or percentiles; the difference between being in the lowest 10% of index values and being in the lowest 20% of index values is 10 percentage points) and of standard deviations.

In the aggregate models, the fixed effect with the largest effect size was focal dominance rank (Table 3). In the aggregate index of aggression given, the relative frequency of aggression given by a focal holding the highest rank position (i.e., the alpha female) was predicted to be 52 percentage points (or 2.7 standard deviations) higher than a focal holding the lowest rank position. In other words, moving from the lowest rank position to the highest rank position is predicted to increase a female’s index value by 52% of the total range of the aggression given index values observed in our population. Conversely, in both of our aggregate indices of submission given (to an aggressive opponent and to a neutral opponent), the relative frequency of submission given by a focal holding the highest rank position was predicted to be 34 percentage points (1.9 s.d.) lower than a focal holding the lowest rank position (Table 3). Age also decreased the relative rate of all agonistic interactions (Table 3). Several other fixed effects had statistically significant relationships with the aggregate and dyadic indices, but these effects were very small (Table 3).

**Table 3.**
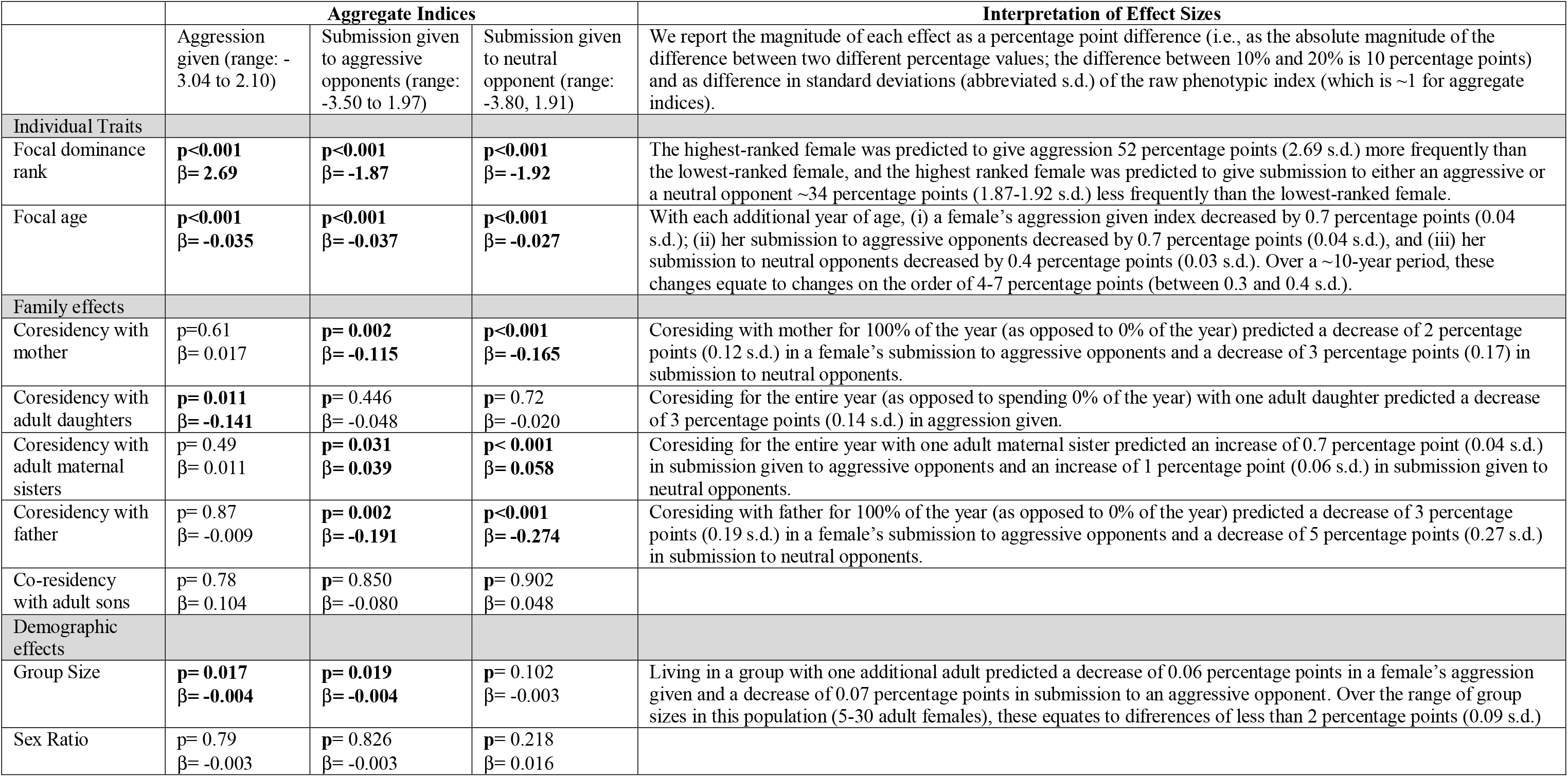
Aggregate models: Effects of fixed predictors on aggregate indices of agonistic interactions. β estimates of effects with p-values are reported for each model. P-values for fixed effects were determined by Wald Conditional Tests.

|  | Aggregate Indices |  |  | Interpretation of Effect Sizes |
|---|---|---|---|---|
|  | Aggression given (range: -3.04 to 2.10) | Submission given to aggressive opponents (range: -3.50 to 1.97) | Submission given to neutral opponent (range: -3.80, 1.91) | We report the magnitude of each effect as a percentage point difference (i.e., as the absolute magnitude of the difference between two different percentage values; the difference between 10% and 20% is 10 percentage points) and as difference in standard deviations (abbreviated s.d.) of the raw phenotypic index (which is ~1 for aggregate indices). |
| Individual Traits |  |  |  |  |
| Focal dominance rank | **p<0.001**<br>**β= 2.69** | **p<0.001**<br>**β= -1.87** | **p<0.001**<br>**β= -1.92** | The highest-ranked female was predicted to give aggression 52 percentage points (2.69 s.d.) more frequently than the lowest-ranked female, and the highest ranked female was predicted to give submission to either an aggressive or a neutral opponent ~34 percentage points (1.87-1.92 s.d.) less frequently than the lowest-ranked female. |
| Focal age | **p<0.001**<br>**β= -0.035** | **p<0.001**<br>**β= -0.037** | **p<0.001**<br>**β= -0.027** | With each additional year of age, (i) a female's aggression given index decreased by 0.7 percentage points (0.04 s.d.); (ii) her submission to aggressive opponents decreased by 0.7 percentage points (0.04 s.d.), and (iii) her submission to neutral opponents decreased by 0.4 percentage points (0.03 s.d.). Over a ~10-year period, these changes equate to changes on the order of 4-7 percentage points (between 0.3 and 0.4 s.d.). |
| Family effects |  |  |  |  |
| Coresidency with mother | p=0.61<br>β= 0.017 | **p= 0.002**<br>**β= -0.115** | **p<0.001**<br>**β= -0.165** | Coresiding with mother for 100% of the year (as opposed to 0% of the year) predicted a decrease of 2 percentage points (0.12 s.d.) in a female's submission to aggressive opponents and a decrease of 3 percentage points (0.17) in submission to neutral opponents. |
| Coresidency with adult daughters | **p= 0.011**<br>**β= -0.141** | **p**= 0.446<br>β= -0.048 | **p**= 0.72<br>β= -0.020 | Coresiding for the entire year (as opposed to spending 0% of the year) with one adult daughter predicted a decrease of 3 percentage points (0.14 s.d.) in aggression given. |
| Coresidency with adult maternal sisters | p= 0.49<br>β= 0.011 | **p= 0.031**<br>**β= 0.039** | **p< 0.001**<br>**β= 0.058** | Coresiding for the entire year with one adult maternal sister predicted an increase of 0.7 percentage point (0.04 s.d.) in submission given to aggressive opponents and an increase of 1 percentage point (0.06 s.d.) in submission given to neutral opponents. |
| Coresidency with father | p= 0.87<br>β= -0.009 | **p= 0.002**<br>**β= -0.191** | **p<0.001**<br>**β= -0.274** | Coresiding with father for 100% of the year (as opposed to 0% of the year) predicted a decrease of 3 percentage points (0.19 s.d.) in a female's submission to aggressive opponents and a decrease of 5 percentage points (0.27 s.d.) in submission to neutral opponents. |
| Co-residency with adult sons | p= 0.78<br>β= 0.104 | **p**= 0.850<br>β= -0.080 | **p**= 0.902<br>β= 0.048 |  |
| Demographic effects |  |  |  |  |
| Group Size | **p= 0.017**<br>**β= -0.004** | **p= 0.019**<br>**β= -0.004** | **p**= 0.102<br>β= -0.003 | Living in a group with one additional adult predicted a decrease of 0.06 percentage points in a female's aggression given and a decrease of 0.07 percentage points in submission to an aggressive opponent. Over the range of group sizes in this population (5-30 adult females), these equates to difrerences of less than 2 percentage points (0.09 s.d.) |
| Sex Ratio | p= 0.79<br>β= -0.003 | **p**= 0.826<br>β= -0.003 | **p**= 0.218<br>β= 0.016 |  |

The dyadic models showed a similar pattern: the largest effect sizes were associated with the focal’s dominance rank, the opponent’s dominance rank, and the interaction between the two (Table 4). Dyads composed of individuals who were close in rank had lower relative frequencies of AS and NS interactions than dyads composed of individuals who were distant in rank (Table 4). As with the aggregate models, several familial and demographic fixed effects were statistically significantly associated with both the AS and the NS indices. Relative rates of agonistic interactions within a dyad decreased with increasing age of either the focal or her opponent, and were lower in mother-daughter and maternal sister dyads (Table 4). All results are reported in Table 4. Using log-transformed dyadic indices did not qualitatively change the fixed effects results (Supplementary Table S8).

**Table 4.** Dyadic models: Effects of fixed predictors on dyadic indices of agonistic interactions. β estimates of effects with p-values are reported for each model. P-values were determined by Wald Conditional Tests.

|  | Dyadic Indices |  | **Interpretation of Effect Sizes** |
|---|---|---|---|
|  | AS (range: -0.008, 0.128) | NS (range: -0.019, 0.370) | We report the magnitude of each effect as a percentage point difference (i.e., as the absolute magnitude of the difference between two different percentage values; the difference between 10% and 20% is 10 percentage points) and as difference in standard deviations (abbreviated s.d.) of the raw phenotypic index. |
| Individual Traits |  |  |  |
| Focal dominance rank | **p<0.001**<br>**β= 0.003** | **p<0.001**<br>**β= -0.029** | In the AS index, the frequency of aggressive gestures for a focal holding the highest rank position was predicted to be 1.8 percentage points (0.3 s.d.) higher than a focal holding the lowest rank position. In the NS index, the frequency of submissive gestures for a focal holding the highest rank position was 7.5 percentage points (1.7 s.d.) lower than a focal holding the lowest rank position. |
| Opponent dominance rank | **p<0.001**<br>**β= -0.014** | **p<0.001**<br>**β= 0.017** | In the AS index, the frequency of aggressive gestures against an opponent holding the highest rank position was predicted to be 10.3 percentage points (1.7 s.d.) lower than against an opponent holding the lowest rank position. In the NS index, the frequency of submissive gestures against an opponent holding the highest rank position was predicted to be 4.3 percentage points (0.9 s.d.) higher than against an opponent holding the lowest rank position. |
| Interaction between focal and opponent dominance rank | **p<0.001**<br>**β= 0.016** | **p<0.001**<br>**β= 0.026** | In the AS index, when the rank difference between focal and the opponent was at the minimum, the AS index was predicted to be 11.9 percentage points (2 s.d.) lower than when the rank difference was at the maximum. When the rank difference between focal and opponent is at the minimum, the NS index was predicted to be 6.6 percentage points (1.4 s.d.) lower than when the rank difference is at its maximum. |
| Focal age | **p<0.001**<br>**β= -4.1e-4** | **p**= 0.128<br>β= -3.8e-4 | In the AS index, when the focal's age increased by one year, the AS index was expected to decrease by 0.3 percentage points (0.05 s.d.). |
| Opponent age | **p<0.001**<br>**β= -2.5e-4** | **p**= 0.587<br>β= -3.4e-4 | In the AS index, when the opponent's age increased by one year, the AS index decreased by 0.2 percentage points (0.03 s.d.). |
| Focal-opponent age interaction | **p<0.001**<br>**β= 1.5e-5** | **p<0.001**<br>**β= 3.2e-5** | When the difference between focal and opponent age increased by one year, AS increased by 0.01 points (< 0.01 s.d.) and NS indices increased by < 0.01 percentage points. |
| Dyad Traits |  |  |  |
| Mother-daughter pair | **p<0.001**<br>**β= -3.1e-3** | **p<0.001**<br>**β= -8.3e-3** | Dyads composed of mother-daughter pairs had AS indices ~ 2.2 percentage points (0.4 s.d.) lower than other dyads and NS indices ~2.1 percentage points (0.5 s.d.) lower than other dyads. |
| Maternal sister pair | **p<0.001**<br>**β= -1.3e-3** | **p<0.001**<br>**β= -3.5e-3** | Dyads composed of maternal sister pairs had AS indices ~ 1 percentage points (0.2 s.d.) lower than other dyads and NS indices ~ 0.9 percentage points (0.2 s.d.) lower than other dyads. |
| Relatedness to opponent | **p**= 0.797<br>β= 2.3e-4 | **p= 0.006**<br>**β= 5.2e-3** | NS interactions occurred more frequently when the individuals involved were more closely related: dyads composed of individuals with 0.5 relatedness (full siblings, mother-daughter dyads) interacted ~ 0.7 percentage points (0.2 s.d.) more than dyad composed of unrelated individuals. |
| Demographic effects |  |  |  |
| Group Size | **p<0.001**<br>**β= -1.6e-4** | **p<0.001**<br>**β= -3.9e-4** | When the group increased in size by 1 adult individual, AS and NS indices both decreased of 0.1 percentage points (0.02 s.d.). Over the typical range of group sizes in this population (5-30 adult females), this equates to differences of 0.5-3 percentage points (0.1-0.6 s.d.). |
| Sex Ratio | **p= 0.004**<br>**β= 2.7e-4** | **p<0.001**<br>**β= 1.4e-3** | A group that is 100% female would have AS and NS indices ~ 0.1-0.2 percentage points (0.02-0.04 s.d.) higher than a group that was 50% female. |

## DISCUSSION

This study is one of only a handful to have investigated the genetic basis of variation in naturally occurring agonistic social interactions in wild vertebrates. We identify measurable heritability in agonistic behaviors in wild female baboons, similar in magnitude to estimates obtained in other animal populations: Lea *et al*. (2010) and Yoshida *et al*. (2016) found heritability values between 0.03 and 0.3 for various naturally occurring agonistic behaviors, with Dochtermann *et al*. (2019) reporting a more general meta-analytic average heritability of aggressive behavior of 0.29 across both vertebrates and invertebrates. Our work places these estimates in context, however, by separately measuring both aggression and submission, and by showing that the underlying quantitative genetics of aggressive and submissive behavior differ in several respects.

For example, based on our aggregate indices, aggressive behavior is approximately twice as heritable as submissive behavior, indicating a proportionally stronger genetic contribution to aggression than submission (Table 1, Figure 2). This result is in line with our findings for the dyadic indices, where an individual’s own genotype explained more variance in aggressive behavior than her opponent’s genotype (i.e., the proportion of variance explained by direct genetic effects was larger than that explained by indirect genetic effects; Table 2, Figure 3). We also found differences in the influence of IGEs on aggressive versus submissive behavior. In particular, in the dyadic NS index, where the focal individual submitted to a neutral opponent, the genotype of her opponent explained more variance than her own genotype (i.e., the contribution of IGEs was larger than that of DGEs). In contrast, the contribution of DGEs was larger than IGEs for the dyadic AS index, which showed a much smaller, although still measurable, IGE (Table 2, Figure 3). In other words, within a given dyad, the genotype of the recipient of unsolicited submissive gestures affected the relative frequency of receiving submission more than the genotype of the individual performing submissive gestures. IGEs rarely account for more variance than DGEs, even for behavioral traits (Santostefano *et al*. (2025); but see one documented case for aggression in crickets: Han *et al*. (2018). However, our result is generally consistent with those of Lea *et al*. (2010), who found that measures of receiving aggressive gestures in yellow-bellied marmots, which they modelled as DGEs, were weakly heritable. Notably, in our data set, the permanent environmental variance (i.e., the non-genetic variance linked to an individual’s identity) also influenced aggression and the receipt of submission more than it influenced submission itself, parallel to our findings for genetic effects.

In addition, we found no genetic correlation, permanent environmental correlation, social status correlation, or phenotypic correlation between a female’s aggressive behavior and her submissive behavior. This result supports the idea that aggression and submission are distinct traits with the potential to evolve independently (Nelson 2005; McCormick & Holekamp 2022). Unfortunately, because our models did not converge when all three covariances (genetic, permanent environmental and social status) were left free to vary, we were not able to fully disentangle genetic, social status, and permanent environmental correlations, limiting our ability to assess their capacity for independent evolution in this population. However, we did find a negative genetic correlation between direct and indirect genetic effects on aggression in our dyadic models, indicating that more aggressive individuals also tend to elicit less aggression from others (as reported, e.g., in Mediterranean field crickets by Santostefano *et al*. (2017). This result implies that the evolution of female aggression in Amboseli baboons is at least partially constrained by a negative correlation between direct and indirect genetic effects, which could contribute to evolutionary stasis for aggressive behavior. That is, even when under selection, aggressive behavior is unlikely to increase in the population because if individuals evolve to become more aggressive, they simultaneously evolve to become less likely to elicit aggression from others.

With respect to submissive behavior, we found a strong, positive genetic correlation between our two aggregate submission indices (r*_a_*=0.96±0.02), indicating that the tendency to be submissive in the face of aggression is correlated with the tendency to be submissive in non-aggressive contexts. Additionally, we identified a negative correlation between DGEs and IGEs for the dyadic NS index, indicating that genotypes that predispose females to give submission to neutral opponents also make those females unlikely to receive unsolicited submissive gestures. Together, these observations suggest that females whose genotype predisposes them to be targeted by aggression also tend to preemptively offer submissive gestures to others, but that they are themselves unlikely to *receive* such unsolicited submissive gestures. These patterns are consistent with a predisposition to “timidity,” where females who are frequently targeted by aggression offer submissive gestures more frequently, possibly to avoid aggression, but are rarely in a position to receive unsolicited submissive gestures themselves. As with the negative genetic correlation between the tendency to be aggressive and the tendency to elicit aggression, the negative correlation between IGEs and DGEs for unsolicited submission may generate some degree of evolutionary constraint on the tendency to be submissive, as selection for an increased frequency of submissive behavior would also select for individuals that tend not to elicit submissive behavior (Santostefano *et al*. 2017).

Several of our conclusions would differ, quantitatively or qualitatively, if we had not accounted for non-genetic sources of social similarity in our model. Failing to account for social inheritance with the **S** matrix led to ∼4-to 5-fold inflated estimates of heritability for the tendency to be submissive to aggressive or neutral opponents. Failing to account for social inheritance also inflated estimates of heritability in aggressive behavior, but to a much smaller extent. This result suggests a relatively stronger influence of genetic variation on variation in aggressive behavior, and a stronger social influence on variation in submissive behavior. Importantly, the effects of the **S** matrix do not simply control for dominance rank itself, which was also included as a fixed effect in all models. Instead, the effects of the **S** matrix indicate that variance in agonistic behavior is also linked to *social similarity*, which is structured by social inheritance of rank in female baboons. This result highlights how the social structure of a society can affect genetic estimates of behavior: moreover, since rank can affect reproductive success and fitness (e.g., Cowlishaw and Dunbar (1991); Alberts *et al*. (2003); Huchard and Lukas (2022), the heritability of fitness itself might also be overestimated if the social variance is conflated with genetic variance.

Not accounting for social inheritance could lead to inflated estimates of heritability in other animal species with complex societies, including humans (Horwitz *et al*. 2003; Young *et al*. 2018; Young *et al*. 2019). Indeed, disentangling the effects of social and genetic inheritance is recognized as one of the most important issues faced by modern human genetics (Xia *et al*. 2021; Young *et al*. 2022), a field where many traits are strongly influenced by social status and wealth, which in turn are frequently influenced by social inheritance (Bowles & Gintis 2002; Piketty 2014; Pfeffer & Killewald 2018). The effects of genotypic variation in humans can now be estimated with family-based genomic methodologies that to a degree account for environmental confounds (Young *et al*. (2018), and these methods can also open the way to investigations of the variance not associated with direct genetic effects (Young *et al*. 2019; Nivard *et al*. 2024; Smith *et al*. 2025). Nevertheless, these solutions are not always applicable: in wild animal populations, genomic information is not always available, and in human populations the most detailed data on fitness and life history traits often come from long-term historical data that lack genomic information (Pettay *et al*. 2005; Moorad & Walling 2017; Young *et al*. 2023). The social matrix may help to disentangle social and genetic effects, and produce more accurate estimates of heritability, by explicitly partitioning the variance associated with social and genetic inheritance and by comparing their effects on the same scale when fine-scale genomic information is not available. Notably, the multi-matrix approach is most useful when there is a strong prior expectation that at least one aspect of the social environment is inherited socially (i.e., not due to genetics, even if genetic and social similarity are correlated), which is well established for female rank in several primates and hyenas (Holekamp & Smale 1991).

The sizable standing genetic variation in aggressive behavior that we found even after accounting for the social environment indicates that considerable genetic variation has been maintained for this trait. This observation may imply that aggressive behavior in female baboons, even if sometimes advantageous, has not been under consistent directional selection in this population (Taylor *et al*. 2014; Kilgour *et al*. 2018) – for example, because nepotistic inheritance of dominance rank in females weakens selection on aggressive behavior. A logical next step therefore involves testing if aggressive behavior is indeed under selection in this population, and, if so, whether it is capable of responding to selection based on a genetic correlation with fitness (Tuliozi *et al*. 2026). Alternatively, the relatively large influence of social status on submissive behavior towards an aggressive opponent suggests that rank strongly determines how much aggression an individual receives, which itself could have strong direct ties to fitness.

Finally, the evolution of female aggression may be driven in part by pleiotropic associations with aggression in male baboons, where rank is not linked to nepotistic social support but almost certainly fitness-relevant (Alberts *et al*. 2003). Understanding which of these explanations apply in this and other wild populations represents an important step forward in explaining the origins and future trajectory of these striking, conspicuous interactions.

## Author contributions

BT: methodology, formal analysis, writing (original draft & review/editing). EMM: conceptualization, methodology, formal analysis, writing (original draft & review/editing), funding acquisition. JT: resources, data curation, writing (review/editing), funding acquisition. EAA: resources, data curation, writing (review/editing), funding acquisition. SCA: conceptualization, supervision, resources, data curation, writing (review/editing), funding acquisition.

## Funding

We gratefully acknowledge the support of the National Science Foundation and the National Institutes of Health for the majority of the data represented here, currently through R01AG071684, R01AG075914, and R3AG078470. Current support for field-based data collection also comes from the Max Planck Institute for Evolutionary Anthropology and Duke University. For a complete set of funding sources, logistical assistance, and data collection and management, please visit http://amboselibaboons.nd.edu/acknowledgements/.

## Conflict of interest statement

The authors declare no conflict of interest.

## Acknowledgments

We thank Duke University, Princeton University, the University of Notre Dame, the Max Planck Institute for Evolutionary Anthropology for financial and logistical support at various times over the years. For assistance and cooperation in Kenya, we are grateful to the Kenya Wildlife Service (KWS), the Wildlife Research Training Institute (WRTI), University of Nairobi, the Kenya Institute of Primate Research (KIPRE), National Museums of Kenya (NMK), National Environment Management Authority (NEMA), and National Commission for Science,Technology, and Innovation (NACOSTI). We thank Jeanne Altmann for her foundational contributions to the Amboseli Baboon Project dataset. We also thank the members of the Amboseli-Longido pastoralist communities, the Enduimet Wildlife Management Area, Tortilis Camp, Ker & Downey Safaris, Serena Lodge, Air Kenya, and Safarilink. Particular thanks go to the current Amboseli Baboon Project long-term data collection team (J.K. Warutere, L. Musembei, D.K. Tajeu), and to T. Wango, and V. Oudu for their assistance in Nairobi. The baboon project database, Babase, is managed by J. Gordon and C. Broderick. For a complete set of acknowledgments please visit http://amboselibaboons.nd.edu/acknowledgements/.

## Notes

### Competing Interest Statement

The authors have declared no competing interest.

