## Supplementary Materials for "Both genetic and social inheritance influence female aggressive and submissive behavior in a wild primate"

### Supplementary Methods S1: Relationship between agonistic behavior frequency and observer effort

The methodology we used here is the same as described in (McLean *et al.* 2023) for the aggregate grooming index, but is applied here to agonistic interactions. From our observations of agonistic behavior, we calculated, for each study subject the relative rate of each type of agonistic behavior (aggression given, submission given to aggressive opponents, or submission given to neutral opponents) she was observed to engage in with any other adult female in the population, by examining daily counts of each interaction type. Because the number of person-hours we devote to observations of each group does not increase with increasing group size, the relative per-capita rate of each agonistic behavior can appear higher in smaller social groups than in larger groups. This difference arises as a simple artifact of having a constant number of observers regardless of group size. Our agonistic interaction data are collected throughout daily observations, during group censuses and the collection of 10-minute focal animal samples, which we conduct by rotating through adult females in random order, completing each rotation before we begin another. Thus, the number of focal animal samples per female per day changes systematically as a function of group size in a manner that reflects the per capita amount of time we spend directly observing each female. The number of focal animal samples per female per day is therefore a good measure of per-female observer effort (see (Archie *et al.* 2014; Campos *et al.* 2020) for additional details).

To correct for the fact that the number of observers does not change with group size, we (separately) regressed daily relative rates of each agonistic behavior for all adult females alive in the population during a given time period against observer effort, measured as the number of focal animal samples collected in a given female’s group per adult female per day. Our agonism indices were the residuals of these regressions. In other words, if we wanted to measure ACA’s aggression given in a 365-day period, we would determine the aggression given and observer effort for all the females alive in our population in that 365-day period. We would use this information to regress the relative rate of aggression given on observer effort. ACA’s residual from that regression (which reflects whether she aggressed other females more or less than the average expected based on observer effort) would be her aggregate index of aggression given for that time period. Note that this methodology both controls for observer effort and standardizes relative individual rates of agonistic behavior across years.

### Supplementary Methods S2: Calculation of aggregate indices of agonistic behavior

To create our aggregate indices of agonistic behavior for each female, for each year she was observed, we regressed her daily rate of each agonistic behavior (aggression given, submission given to aggressive opponents or submission given to neutral opponents) in that year on observer effort (see Supplementary Methods S1). This annual mean value was not based on the calendar year, but on each female’s birth date and hence age class (e.g., her 6th year of age, 7th year of age, and so on). This approach is computationally expensive. It means that, for example, if we wanted to measure a baboon’s aggression given between her fifth and sixth birthday, we would determine the aggression given frequency and observer effort for all the females alive in our population in that specific 365-day period. We would use these data to regress aggression given frequency on observer effort. The only data point we would use from this regression is the residual for our one specific baboon of interest. We calculated the indices in this manner for each adult female in each year of her life in which she had at least 60 days of observations during the study period (January 1983-June 2017); these included years in which a female reached maturity or died partway through the year. Note that the average index value is close to 0 – females who participated in more interactions than average have higher, positive values and females who participated in fewer interactions than average have lower, negative values.

### Supplementary Methods S3: Calculation of dyadic indices of agonistic behavior

To explicitly investigate the role of indirect genetic effects on agonistic behavior, we calculated (separately for aggressive-submissive AS and neutral-submissive NS interactions) for each pair of adult females, for each year that they were co-resident in a social group for at least 60 days in that year and that had at least one agonistic interaction of that kind. We used the same set of females and female years as in the aggregate indices. However, for ease of calculation, and because females in a dyad had different birthdates, we calculated the dyadic index for each calendar year instead of for each year of age for a given female. The dyadic indices were based on the daily relative rate of agonistic interactions involving the focal individual and a specific opponent over the course of a calendar year, considering only the days where both focal and opponent were present as adults in the same social group, and thereby available to interact with each other.

As with the aggregate indices described above, the dyadic index is the residual of the regression of this daily relative agonistic interaction rate on observer effort in the year of interest. Positive values indicate dyads in which the focal gave above average amounts of aggression (AS index) or submission (NS index) to that particular opponent (relative to the mean of the population, controlling for observer effort) and negative values indicate dyads in which the focal gave below average amounts of aggression (AS index) or submission (NS index) to that particular opponent (controlling for observer effort) as compared to the entire population. Dyadic agonistic indices do not have to be (and overwhelmingly are not) symmetrical between the two females in a dyad: for approximately 95% of the dyad-years in our study, agonistic interactions occurred in only one direction, i.e., individual 1 only gave aggressive gestures and individual 2 only gave submissive gestures. Thus, for 95% of dyads, one of the two yearly values for the AS (and NS) index was derived from a count of 0 interactions.

### Supplementary Results S4: Genetic ancestry

The baboons roaming the Amboseli basin represent an admixed population, with admixture resulting from both historical and recent admixture events. Most individuals have majority yellow baboon ancestry (*Papio cynocephalus*) with varying amounts of admixed anubis baboon ancestry (*P. anubis*) (Wall *et al.* 2016; Vilgalys *et al.* 2022). These two species show little evidence of ecological differentiation (Winder 2015; Wango *et al.* 2019) and produce viable and fertile offspring with little evidence of hybrid dysgenesis (Ackermann *et al.* 2006; Tung *et al.* 2008). However, molecular genetic analysis has revealed selection against admixture in the study population (Vilgalys *et al.* 2022) and genetic ancestry is associated with variation in several behavioral and fitness-related traits (Charpentier *et al.* 2008; Tung *et al.* 2012; Franz *et al.* 2015; Fogel *et al.* 2021). We did not include anubis ancestry as an effect in our main models because we lacked estimates of genetic ancestry for 200 of our study subjects (55%). Thus, to investigate whether hybrid ancestry influenced how variance was partitioned in our analyses of agonistic behavior, we first identified the subset of 165 subjects for whom we ancestry estimates were available, and then re-ran the same animal models reported in the main text using only this subset of subjects. We fit these models both with and without ancestry estimates (represented as the proportion of introgressed anubis ancestry in each individual’s genome, from Vilgalys *et al.* (2022) included as a fixed effect).

All parameters estimated by the models that included the fixed effect of introgressed anubis ancestry were very similar to those estimated by the models that excluded the ancestry parameter (Supplementary Table S4). Accordingly, genetic ancestry scores do not appear to influence our estimates of trait heritability, genetic correlation between agonistic behaviors, or DGEs and IGEs for agonistic behaviors. Aggregate submission given to neutral opponents did decrease slightly with increased anubis ancestry (p = 0.034, β = -0.810), but including ancestry estimates did not change the significance and direction of other fixed effects. While the reduced sample size limits the scope of our interpretation, these results suggest that while overall introgressed anubis ancestry does not have a detectable association with agonistic behavior, individuals with more anubis ancestry may be slightly less likely to give submission to neutral opponents.

### Supplementary Methods S5: Linear forms of all aggregate and dyadic models (all nested steps)

Aggregate models

Null model:

$y_{ik}=fixed effects+e_{ik}$	(1)

Repeatability model:

$y_{ik}=fixed effects+F_{i}+e_{ik}$	(2)

Heritability model:

$y_{ik}=fixed effects+a_{i}+{pe}_{i}+e_{ik}$	(3)

Heritability & status value model:

$y_{ik}=fixed effects+a_{i}+{pe}_{i}+s_{i}+e_{ik}$	(4)

In these models, $y_{ik}$ is the aggregate agonism index for (focal) individual *i* in year *k*, $e_{ik}$ is a residual error term, $F_{i}$ is a random effect of the identity of the focal individual, $a_{i}$is the additive genetic contribution of individual *i* (i.e., her breeding value; note that since individual *i* is the focal, this contribution could also be written as $a_{F_{i}}$), ${pe}_{i}$ represents the permanent environment effect of individual *i*, and $s_{i}$ is the social status contribution of individual *i* (its contribution to the **S**-matrix).

Dyadic models

Null model:

$y_{ijk}=fixed effects+{dyad}_{ij}+e_{ijk}$	(5)

Repeatability model:

$y_{ijk}=fixed effects+F_{i}+{dyad}_{ij}+e_{ijk}$	(6)

Repeatability with opponent model:

$y_{ijk}=fixed effects+F_{i}+O_{j}+{dyad}_{ij}+e_{ijk}$	(7)

Direct heritability model (with direct genetic effect (DGE)):

$y_{ijk}=fixed effects+a_{F_{i}}+{pe}_{F_{i}}+O_{j}+{dyad}_{ij}+e_{ijk}$	(8)

Dyadic heritability model (DGE + IGE):

$y_{ijk}=fixed effects+a_{F_{i}}+{pe}_{F_{i}}+a_{O_{j}}+{pe}_{O_{j}}+{dyad}_{ij}+e_{ijk}$	(9)

Direct heritability & direct status model (DGE + Direct status value (DSE)):

$y_{ijk}=fixed effects+a_{F_{i}}+{pe}_{F_{i}}+S_{F_{i}}+O_{j}+{dyad}_{ij}+e_{ijk}$	(9.1)

Dyadic heritability and direct status model (DGE + DSE + indirect genetic effect (IGE)):

$y_{ijk}=fixed effects+a_{F_{i}}+{pe}_{F_{i}}+S_{F_{i}}+a_{O_{j}}+{pe}_{O_{j}}+{dyad}_{ij}+e_{ijk}$	(9.2)

Dyadic heritability model (DGE + IGE) with added status values of both focal (DSE) and opponent (indirect social effect ISE):

$y_{ijk}=fixed effects+a_{F_{i}}+{pe}_{F_{i}}+S_{F_{i}}+a_{O_{j}}+{pe}_{O_{j}}+S_{O_{j}}+{dyad}_{ij}+e_{ijk}$	(10)

In these models, $y_{ijk}$ is the frequency of interactions between focal individual *i* and opponent individual *j* in year *k.* In the “AS” index, the focal individual *i* gave aggressive gestures, while in the “NS” index the focal individual *i* gave submissive gestures. ${dyad}_{ij}$is an identity assigned to each unique pair of individuals; this term is included because we have repeated measures for each dyad in the dataset. As with the aggregate index models, $e_{ijk}$ is residual error, $F_{i}$ is the random effect of the focal’s identity, $a_{F_{i}}$is the additive genetic contribution of the focal *i*, ${pe}_{F_{i}}$ is the permanent environment effect of the focal, and $S_{F_{i}}$ is the random effect of individual *i*’s social status. Note that in dyadic indices the terms with subscript $i$all refer to the focal. In addition, $O_{j}$ is a random effect of the identity of the opponent (who is performing submissive gestures in the AS dyadic index, and neutral or no gestures in the NS dyadic index), $a_{O_{j}}$ is the additive genetic contribution of the opponent, ${pe}_{O_{j}}$ is the permanent environmental effect of the opponent, and $S_{O_{j}}$is the social contribution of the opponent.

Because direct and indirect genetic effects ($a_{F_{i}}$ and $a_{O_{j}}$) were free to covary in this best-fitting model, we estimated the direct genetic variance ($\text{σ}_{a_{F_{i}}}^{\text{2}}$), the indirect genetic variance ($\text{σ}_{a_{O_{j}}}^{\text{2}}$) and the covariance between them ($\text{σ}_{a_{F_{i}}a_{O_{j}}}$). The variance-covariance matrix (**G**) for the additive genetic effects with covariances free to vary is as follows, with **A** representing the additive genetic relatedness matrix:

$\mathbf{G}\otimes\mathbf{A}$	(11)

$\mathbf{G}=\left| \begin{matrix} \text{σ}_{a_{F_{i}}}^{\text{2}} & \text{σ}_{a_{F_{i}}a_{O_{j}}} \\ \text{σ}_{{a_{O_{j}}a}_{F_{i}}} & \text{σ}_{a_{O_{j}}}^{\text{2}} \end{matrix} \right|$	(12)

In the best-fitting model for dyadic aggression index, permanent environmental covariance was also left free to covary, with permanent environmental variance-covariance matrix **PE**:

$\mathbf{PE}=\left| \begin{matrix} \text{σ}_{{PE}_{F_{i}}}^{\text{2}} & \text{σ}_{{PE}_{F_{i}}{PE}_{O_{j}}} \\ \text{σ}_{{{PE}_{O_{j}}PE}_{F_{i}}} & \text{σ}_{{PE}_{O_{j}}}^{\text{2}} \end{matrix} \right|$	(12)

### Supplementary Methods S6: Variance-covariance structure of all bivariate models of aggregate agonism indices

We built two bivariate models using aggregate indices: 1) between aggression given and submission given to an aggressive opponent; and 2) between submission given to an aggressive opponent and submission given to a neutral opponent. For each of these two bivariate analyses, we built four combinations of variance-covariance structures. Trait 1 (T1) and Trait 2 (T2) represent the two traits included in all bivariate analysis, such that $\text{σ}_{a_{T1}}^{\text{2}}$, $\text{σ}_{a_{T2}}^{\text{2}}$, and $\text{σ}_{a_{T2}a_{T1}}$ represent the genetic variance of Trait 1, Trait 2 and the genetic covariance between the two, respectively. Similarly, $\text{σ}_{{PE}_{T1}}^{\text{2}}$, $\text{σ}_{{PE}_{T2}}^{\text{2}}$, and $\text{σ}_{{PE}_{T2}{PE}_{T1}}$ represent the permanent environmental variance of Trait 1, Trait 2 and the covariance between the two, respectively; and $\text{σ}_{S_{T1}}^{\text{2}}$, $\text{σ}_{S_{T2}}^{\text{2}}$, and $\text{σ}_{S_{T2}S_{T1}}$ represent the social status variance of Trait 1, Trait 2 and the covariance between the two, respectively. Matrix **A** is the additive genetic relationship matrix, **I** is an identity matrix of appropriate size, and **S** is the social status matrix. The model structures are defined below.

**Bivariate model structure 1**. Genetic, PE and Social covariances constrained to zero.

For the genetic variance-covariance matrix $\mathbf{G}$:

$\mathbf{G}\otimes\mathbf{A}$	(BIV1)

$$\mathbf{G}=\left| \begin{matrix} \text{σ}_{a_{T1}}^{\text{2}} & \text{0} \\ \text{0} & \text{σ}_{a_{T2}}^{\text{2}} \end{matrix} \right|$$

For the permanent environmental variance-covariance matrix $\mathbf{PE}$:

$\mathbf{PE}\otimes\mathbf{I}$	(BIV2)

$$\mathbf{PE}=\left| \begin{matrix} \text{σ}_{{PE}_{T1}}^{\text{2}} & \text{0} \\ \text{0} & \text{σ}_{{PE}_{T2}}^{\text{2}} \end{matrix} \right|$$

For the social variance-covariance matrix $\mathbf{ST}$:

$\mathbf{ST}\otimes\mathbf{S}$	(BIV3)

$$\mathbf{ST}=\left| \begin{matrix} \text{σ}_{S_{T1}}^{\text{2}} & \text{0} \\ \text{0} & \text{σ}_{S_{T2}}^{\text{2}} \end{matrix} \right|$$

**Bivariate model structure 2**. Social covariance allowed to vary; genetic and permanent environmental covariance constrained to zero.

The genetic variance-covariance matrix is as indicated in eq. BIV1. The permanent environmental variance-covariance matrix is as indicated in eq. BIV2. For the social status variance-covariance matrix $\mathbf{ST}$:

$\mathbf{ST}\otimes\mathbf{S}$	(BIV4)

$$\mathbf{ST}=\left| \begin{matrix} \text{σ}_{S_{T1}}^{\text{2}} & \text{σ}_{S_{T1}S_{T2}} \\ \text{σ}_{S_{T2}S_{T1}} & \text{σ}_{S_{T2}}^{\text{2}} \end{matrix} \right|$$

**Bivariate model structure 3**. Genetic covariance allowed to vary; social status and permanent environmental covariance constrained to zero.

The permanent environmental variance-covariance matrix is as indicated in equation BIV2 and the social status variance-covariance matrix is as indicated in eq. BIV3. For the genetic variance-covariance matrix $\mathbf{G}$:

$\mathbf{G}\otimes\mathbf{A}$	(BIV5)

$$\mathbf{G}=\left| \begin{matrix} \text{σ}_{a_{T1}}^{\text{2}} & \text{σ}_{a_{T1}a_{T2}} \\ \text{σ}_{a_{T2}a_{T1}} & \text{σ}_{a_{T2}}^{\text{2}} \end{matrix} \right|$$

**Bivariate model structure 4**. Permanent environmental covariance allowed to vary; social status and genetic covariance constrained to zero.

The genetic variance-covariance matrix is as indicate in eq. BIV1. The social status variance-covariance matrix is as indicated in equation BIV3. For the permanent environmental variance-covariance matrix $\mathbf{PE}$:

$\mathbf{PE}\otimes\mathbf{I}$	(BIV6)

$$\mathbf{PE}=\left| \begin{matrix} \text{σ}_{{PE}_{T1}}^{\text{2}} & \text{σ}_{{PE}_{T1}{PE}_{T2}} \\ \text{σ}_{{PE}_{T2}{PE}_{T1}} & \text{σ}_{{PE}_{T2}}^{\text{2}} \end{matrix} \right|$$

All bivariate models where more than one unstructured variance-covariance was fitted did not converge – in other words, all bivariate models converged only when at least two covariances out of three (except the residual) were constrained to 0. This restricts the scope of our discussion, and it was likely because limited power created identifiability problems and because several correlations were low and non-significant.

### Supplementary Results S7: The role of the social status matrix in dyadic phenotypes

In our population, including the social status matrix improved the models for our aggregate phenotypes (Table 1, main text), but apparently did not improve the models for any of our dyadic phenotypes (Table 2). Thus, the social inheritance of dominance rank explains variance in aggregate measures of aggression and submission, but it might play less of a role in shaping variation in the frequency of interactions within specific dyads of individuals. In other words, while individuals with similar social status engage in agonistic behaviors at similar frequencies overall, the way individuals distributed that behavior among social opponents is not clearly structured by similarity in their social status. For example, some individuals who occupy a particular lifetime average dominance rank may distribute their aggression in a relatively concentrated manner, focusing it on few individuals. On the other hand, other individuals with similar dominance rank may distribute their aggression more widely to several social opponents, depending on their web of specific relationships.

Individual- and dyad-level variation of this nature means that some of the variance in the aggregate models that was associated with the random effect of social status would be absorbed, in the dyadic models, by opponent identity (indirect genetic effects and permanent environment) and dyad identity (dyad random term). This explanation would also make it more likely for variance that was associated with similarity in social status in the aggregate model to be associated with the random effect of “dyad” in the dyadic models. This interpretation is in line with the view that submissive behavior is plastic: individuals with similar social status will behave similarly overall, but they will adjust their submissive behavior according to different social contexts (i.e. their opponent within a given dyad). Finally, note that variance estimates are smaller in dyadic indices than in their aggregate counterparts: this difference in magnitude likely arises from the fact that any given dyad in the dyadic dataset has many fewer interactions than any given focal individual in the aggregate dataset. As a result, small errors in measurement have a larger effect on our dyadic index than on our aggregate index, inflating the residual (error) variance in our dyadic index and producing a conservative estimate of the genetic effects for both members of the dyad.

Overall, these results indicate that, whenever the individuals involved in the interactions are known, using both a multi-matrix and an indirect genetic effects approach can offer complementary information on the nature of social and genetic influence on behavior.

### Supplementary Tables

#### **Table S1**. Behaviors that constitute agonistic interactions and their descriptions^1^

| **Behavior** | **Description** |
| --- | --- |
| **Aggressive behaviors** | |
| Raised brow or eyelid display | Display of unpigmented skin beneath eyebrows either by raising brow itself or tilting head dorsally; often accompanied by jutting forward of the head. |
| Open mouth face or attempted bite | Jaw held open, teeth not exposed or only slightly exposed, often accompanied by forward jutting of the head. |
| Bobbed head and thorax | Abrupt, rapid raising and lowering of head and trunk; body may show a forward movement component. |
| Ground slapped | Palm of hand or hands struck against ground, often audible. |
| Chomped or gave exaggerated chewing motions | Repeated and exaggerated chewing or grinding movements of the jaws, often with extreme lateral excursion, copious salivation and audible grinding. |
| Lunged at | Forward leaping or jumping towards another individual; no lateral movement. |
| Gave threat yawn or directed gape | Directed gape or yawn-like moving, usually exposing canines. |
| Rubbed muzzle on substrate | Muzzle and chin rubbed laterally on ground or tree limbs. Often accompanied by gaping and chomping. |
| Hit or slapped | Rapid, open-handed striking or attempted striking of another individual. |
| Pushed | Open-handed shoving or attempted shoving of another individual. |
| Grabbed | Closed-handed gripping of another individual, usually brief. |
| Held down | Closed-handed gripping of another individual, forcing that individual against substrate, usually prolonged. |
| Bite | Obvious. |
| **Submissive Behaviors** | |
| Averted head and/or body | Turned head aside at approach of another individual, often accompanied by fixedly looking away from the other individual, but without the tensing or twisting associated with a cower. |
| Cowered or leaned aside | Lateral flexion of the spine, often from a seated or crouching position; limbs usually held against the body but not always. |
| Gave fear paralysis | Subject leaned forward or lay on ground in rigid, tensed and motionless position. Often accompanied by screaming, urination and defecation. |
| Grimace | Corners of mouth retracted and teeth (often clenched) exposed, grimace clearly directed towards another individual. |
| Gave tail-up | Tail held upright, or nearly so, or even 'pointed' forward over back (accompanied by other submissive gestures). |
| Gave high-pitched shrill scream or screech | Obvious. |
| Gave cackle or gecker vocalization | Obvious. |
| Repeated rapid glances | Individual glances repeatedly and rapidly in the direction of a conspecific. Sometimes this involves looking over the shoulder or behind; it almost always interrupts the current activity at least briefly. |

^1^Table modified from Alberts *et al.* (2011).

#### **Table S2**. Trimmed pedigree size and structure. The ‘trimmed’ pedigree is the pedigree describing all known genetic relationships between individuals in the dataset. Some paternal half-siblings and full siblings in our dataset may be undetected.

| **Pedigree** | |
| --- | --- |
| Individuals in the dataset | 365 |
| Individuals in the pedigree | 473 |
| Mother-offspring pairs | 389 |
| Father-offspring pairs | 266 |
| Maternal half-siblings pairs | 378 |
| Paternal half-siblings pairs | 392 |
| Full siblings pair | 28 |
| Maximum generation depth | 8 |

#### **Table S3**. Fixed effects included in all models investigating aggregate indices of agonistic behavior.

| **Aggregate indices** | | |
| --- | --- | --- |
| **Fixed effect** | **Definition** | **Reason for inclusion** |
| Individual Traits |  |  |
| Focal dominance rank | Linear effect of proportional rank of the focal, averaged over the index year | This effect tests the linear influence of rank on the phenotype regardless of individual identity, genetic, or social relationships. |
| Focal age | Linear effect of age of the focal in the index year | Age affects multiple aspects of behavior in long-lived mammals and is included for completeness. |
| Family effects |  |  |
| Co-residency with mother | Linear effect of the proportion of time the focal spent co-residing with her mother during the index year | The presence of the mother increases the potential for kin support during conflicts, which can influence rates and outcomes of agonistic interactions (Lea *et al.* 2014). |
| Co-residency with adult daughters | Linear effect of the sum of the proportions of times spent co-residing with each adult daughter in the focal’s group during the index year. If, e.g., a focal individual co-resided with two adult daughters for the full index year, and with a third daughter for only six months during that index year, her value would be 2.5 for this fixed effect. | The higher the number of daughters, the greater the potential for kin support during conflicts, which in turn can influence rates and outcomes of agonistic interactions (Lea *et al.* 2014). |
| Co-residency with adult maternal sisters | Linear effect of the sum of the proportions of times spent with each adult maternal sisters in the focal’s group during the index year. | The higher the number of sisters, the greater the potential for kin support during conflicts, which in turn can influence rates and outcomes of agonistic interactions (Lea *et al.* 2014). |
| Co-residency with father | Linear effect of the proportion of time spent co-residing with the father of the focal in the group during the index year | The presence of the father increases the potential for kin support during conflicts, which in turn can potentially influence rates and outcomes of agonistic interactions (Buchan *et al.* 2003). |
| Co-residency with adult sons | Linear effect of the sum of the proportions of times spent co-residing with each adult son in the focal’s group during the index year. | The presence of a son increases the potential for kin support during conflicts, which in turn can potentially influence rates of agonistic interactions (Engh *et al.* 2009). |
| Demographic effects |  |  |
| Group Size | Linear effect of the number of adult females in the group of the focal, averaged across the index year. The number of adult females is well-correlated with the total number of adults in the group, and with total group size. | Group size can influence the attainment of dominance rank, and may therefore affect rates and outcomes of agonistic interactions (Lea *et al.* 2014). In addition, group size predicts the per capita frequency of agonistic interactions in social groups; individuals in larger groups engage in more agonistic interactions (Lerch et al., in prep). |
| Sex Ratio | Linear effect of the adult sex ratio in the group of the focal, averaged across the index year. | The adult sex ratio in a group can influence competition for mates, and could therefore potentially affect rates of agonistic interactions (but no evidence of this effect was found in Baniel *et al.* (2018)). |

#### **Table S4**. Fixed effects included in all models investigating dyadic indices of agonistic behavior.

| **Dyadic indices** | | |
| --- | --- | --- |
| Fixed effect | Definition | Reason for inclusion |
| Individual Traits |  |  |
| Focal dominance rank | Linear effect of proportional rank of the focal, averaged over the index year | See Table 1 |
| Opponent dominance rank | Linear effect of proportional rank of the opponent, averaged over the index year | See Table 1 |
| Rank interaction | Interaction between the focal dominance rank and opponent dominance rank | Relative interaction frequencies may vary depending upon how close in rank the opponents are (e.g., high-ranking focals may aggress low-ranking opponents more often than they do high-ranking ones). |
| Focal age | Linear effect of age of the focal in the index year | See Table 1 |
| Opponent age | Linear effect of age of the opponent in the index year | See Table 1 |
| Focal-opponent age interaction | Interaction between the focal age and opponent age | The relative interaction frequencies may vary depending upon how close in age the opponents are (e.g., older individuals might aggress older opponents more often than younger opponents). |
| Dyad Traits |  |  |
| Mother-daughter pair | Binary effect of the dyad being a mother-daughter pair | Females’ strongest social bonds are with their mothers, and these bonds may influence the tendency to engage in agonistic interactions (Silk *et al.* 2006a; Silk *et al.* 2006b). |
| Maternal sister pair | Binary effect of the dyad being a mother-sister pair | After mothers, maternal sisters exhibit the strongest social bonds, and these bonds may influence the tendency to engage in agonistic interactions (Silk *et al.* 2006a; Silk *et al.* 2006b). |
| Relatedness to opponent | Linear effect of relatedness with the opponent. | Some primates, including the baboons we studied here, show a graded patterns of social bond strength that is strongest with mother-daughter pairs and decreases with decreasing relatedness, with consequences for the tendency to engage in agonistic interactions (Chapais & Berman 2004; Silk *et al.* 2006b). |
| Demographic effects |  |  |
| Group Size | Linear effect of the number of adult females in the group of the dyad, averaged across the index year | Group size may be associated with density-dependent competition, which in turn could influence the rate of agonistic interactions; dyadic relationships involve more agonisms in larger groups (Lerch et al., in prep) |
| Sex Ratio | Linear effect of the adult sex ratio in the group of the dyad, averaged across the index year | The adult sex ratio of a group could influence competition for mates, and thus potentially the rate of agonistic interactions (but see Baniel *et al.* (2018)) |

#### **Table S5**. Comparisons of models obtained with and without the focal’s estimated genetic ancestry (i.e., the estimated proportion of their genome that represents introgressed anubis ancestry) as a fixed effect.

|  | Model without genetic ancestry, with reduced sample size^2^ | Model with genetic ancestry, with reduced sample size^2^ |
| --- | --- | --- |
| Aggregate aggression given: heritability and status value model | | |
| $\sigma_{a}^{2}$ | 0.083 (0.027) | 0.084 (0.027) |
| $\sigma_{PE}^{2}$ | 0.002 (0.019) | 0.002 (0.019) |
| $\sigma_{S}^{2}$ | 0.013 (0.013) | 0.013 (0.013) |
| $\sigma_{e}^{2}$ | 0.210 (0.008 | 0.210 (0.008) |
| $h^{2}$ | 0.270 (0.078) | 0.270 (0.080) |
| $s^{2}$ | 0.042 (0.040) | 0.041 (0.041) |
| Aggregate submission given to aggressive opponent: heritability and status value model | | |
| $\sigma_{a}^{2}$ | 0.047 (0.016) | 0.041 (0.016) |
| $\sigma_{PE}^{2}$ | <0.001 (NA) | <0.001 (NA) |
| $\sigma_{S}^{2}$ | 0.110 (0.026) | 0.110 (0.026) |
| $\sigma_{e}^{2}$ | 0.267 (0.010) | 0.267 (0.001) |
| $h^{2}$ | 0.100 (0.040) | 0.100 (0.040) |
| $s^{2}$ | 0.267 (0.052) | 0.261 (0.052) |
| Aggregate submission given to neutral opponent: heritability and status value model | | |
| $\sigma_{a}^{2}$ | 0.029 (0.014) | 0.031 (0.014) |
| $\sigma_{PE}^{2}$ | <0.001 (NA) | <0.001 (NA) |
| $\sigma_{S}^{2}$ | 0.168 (0.033) | 0.157 (0.032) |
| $\sigma_{e}^{2}$ | 0.213 (0.008) | 0.213 (0.008) |
| $h^{2}$ | 0.070 (0.035) | 0.077 (0.037) |
| $s^{2}$ | 0.411 (0.056) | 0.391 (0.057) |
| Dyadic AS: dyadic heritability (DGE + IGE) model | | |
| $\sigma_{a_{F}}^{2}$ | 4.6e-6 (7.4e-7) | 4.5e-6 (1.28e-6) |
| $\sigma_{a_{O}}^{2}$ | 1.4e-6 (8.02e-7) | 1.45e-6 (8.22 e-7) |
| $\sigma_{{PE}_{F}}^{2}$ | 1.6e-11 (NA) | 3.87e-8 (7.79e-7) |
| $\sigma_{{PE}_{O}}^{2}$ | 1.23e-6 (6.67e-7) | 1.24e-6 (6.79e-7) |
| $\sigma_{dyad}^{2}$ | 1.05e-6 (2.95e-7) | 1.05e-6 (2.96e-7) |
| $\sigma_{e}^{2}$ | 3.74e-5 (6.13e-7) | 3.74e-5 (6.13e-7) |
| DGE | 0.10 (0.010) | 0.10 (0.010) |
| IGE | 0.03 (0.012) | 0.03 (0.018) |
| Dyadic NS: dyadic heritability (DGE + IGE) model | | |
| $\sigma_{a}^{2}$ | 2.76e-5 (6.76e-6) | 2.6e-5 (6.7e-6) |
| $\sigma_{a_{O}}^{2}$ | 8.4e-6 (3.98e-6) | 8.3e-6 (4e-6) |
| $\sigma_{{PE}_{F}}^{2}$ | 1.08e-6 (4.01e-6) | 2.2e-6 (4.1e-6) |
| $\sigma_{{PE}_{O}}^{2}$ | 4.64e-6 (2.97e-6) | 4.7e-6 (3e-6) |
| $\sigma_{dyad}^{2}$ | 1.05e-5 (1.52e-6) | 1.05e-5 (1.52e-6) |
| $\sigma_{e}^{2}$ | 1.63e-4 (2.76e-6) | 1.63e-4 (2.76e-6) |
| DGE | 0.12 (0.030) | 0.12 (0.030) |
| IGE | 0.04 (0.020) | 0.03 (0.020) |

^2^These models were run with the subset of subjects for whom genetic ancestry was measured (165 out of 365 subjects). All results are quantitatively similar, and in some cases identical, between the two models.

#### **Table S6**. **Correlations from bivariate aggregate models**. We fitted four bivariate models for each phenotype combination (aggression versus submission to aggressive opponent; submission to aggressive opponent versus submission to neutral opponent). In the first three models, we fitted all variance components (genetic, social status, permanent environment) that we used in the main models (Table 1), and we left only one covariance free to vary in each model (either genetic covariance, social status covariance, or permanent environmental covariance, always in addition to the residual covariance); the other covariances were set to zero. In the fourth model (the individual identity model, or repeatability model) we only included the random effect of individual identity, without further partitioning its variance into its genetic, social, and permanent environmental components. Each of the first three models produced an estimate of the phenotypic correlation (last column in the table) as well as an estimate of the genetic correlation or the social status correlation or the permanent environment correlation, depending on which covariance was left free to vary (next-to-last column in the table). The fourth model also produced an estimate of the phenotypic correlation, and an estimate of the “individual identity” correlation, estimated using only individual as a random effect. Significant individual-level correlations are **bolded**.

| Phenotype combinations | Individual-level random effects | Type of covariance free to vary | Correlation | Phenotypic Correlation |
| --- | --- | --- | --- | --- |
| Aggression and Submission to aggressive opponent | Genetic + Social status + Permanent environment | Genetic | 0.23 ± 0.15 | 0.09 ± 0.02 |
|  |  | Social status | 0.25 ± 0.16 | 0.10 ± 0.03 |
|  |  | Permanent environment | 0.47 ± 0.65 | 0.07 ± 0.02 |
|  | Individual identity | Individual identity | 0.14 ± 0.07 | 0.11 ± 0.03 |
| Submission to aggressive opponent and submission to neutral opponent | Genetic + Social status + Permanent environment | Genetic | **0.96 ± 0.01** | 0.69 ± 0.02 |
|  |  | Social status | **0.99 ± 0.01** | 0.65 ± 0.02 |
|  |  | Permanent environment | did not converge | |
|  | Individual identity | Individual Identity | **0.93 ± 0.01** | 0.70 ± 0.01 |

**Table S7**. Variances of dyadic models run with log-transformed AS and NS dyadic agonism indices, showing estimates of variance components, with standard errors in parentheses below each estimate. Only the best-fitting models from Table 2 are shown.

|  | Model name | $\sigma_{a_{F}}^{2}$ | $\sigma_{a_{O}}^{2}$ | $\sigma_{{PE}_{F}}^{2}$ | $\sigma_{{PE}_{O}}^{2}$ | $\sigma_{dyad}^{2}$ | $\sigma_{e}^{2}$ | DGE | IGE | $\text{r}_{a_{F}a_{O}}$ | $\text{r}_{{PE}_{F}{PE}_{O}}$ |
| --- | --- | --- | --- | --- | --- | --- | --- | --- | --- | --- | --- |
| AS | Dyadic heritability (DGE & IGE) with $\text{r}_{a_{F}a_{O}}$ and $\text{r}_{{PE}_{F}{PE}_{O}}$ | 0.035 (0.009) | 0.015 (0.004) | 0.014 (0.006) | 0.005 (0.003) | 0.008 (0.002) | 0.29 (0.003) | 0.10 (0.02) | 0.04 (0.01) | -0.18 (0.20) | 0.85  (0.34) |
| NS | Dyadic heritability (DGE & IGE) with $\text{r}_{a_{F}a_{O}}$ | 0.011  (0.004) | 0.022  (0.005) | 0.009  (0.003) | 0.006  (0.003) | 0.008  (0.001) | 0.16  (0.002) | 0.053  (0.016) | 0.103  (0.022) | -0.293  (0.129) | - |

**Table S8**. Fixed effects estimates of dyadic models run with log-transformed AS and NS dyadic indices: Effects of fixed predictors on dyadic indices of agonistic interactions. β estimates of effects with p-values are reported for each model. P-values were determined by Wald Conditional Tests.

|  | Dyadic Indices | |
| --- | --- | --- |
|  | AS (range:  -10.405, -1.994) | NS (range  -7.616, -0.943) |
| Individual Traits |  |  |
| Focal dominance rank | **p<0.001**  **β= 0.186** | **p<0.001**  **β= -1.267** |
| Opponent dominance rank | **p<0.001**  **β= -1.369** | **p<0.001**  **β= 0.516** |
| Interaction between focal and opponent dominance rank | **p<0.001**  **β= 1.568** | **p<0.001**  **β= 1.203** |
| Focal age | **p<0.001**  **β= -0.029** | p= 0.119  β= -0.013 |
| Opponent age | **p<0.001**  **β= -0.017** | p= 0.482  β= -0.010 |
| Focal-opponent age interaction | **p<0.001**  **β= 0.001** | **p<0.001**  **β= 0.001** |
| Dyad Traits |  |  |
| Mother-daughter pair | **p<0.001**  **β= -0.247** | **p<0.001**  **β= -0.243** |
| Maternal sister pair | **p<0.001**  **β= -0.104** | **p<0.001**  **β= -0.123** |
| Relatedness to opponent | p= 0.441  β= -0.055 | p= 0.236  β= 0.067 |
| Demographic effects |  |  |
| Group Size | **p<0.001**  **β= -0.003** | **p<0.001**  **β= -0.007** |
| Sex Ratio | **p<0.001**  **β= 0.042** | **p<0.001**  **β= 0.025** |

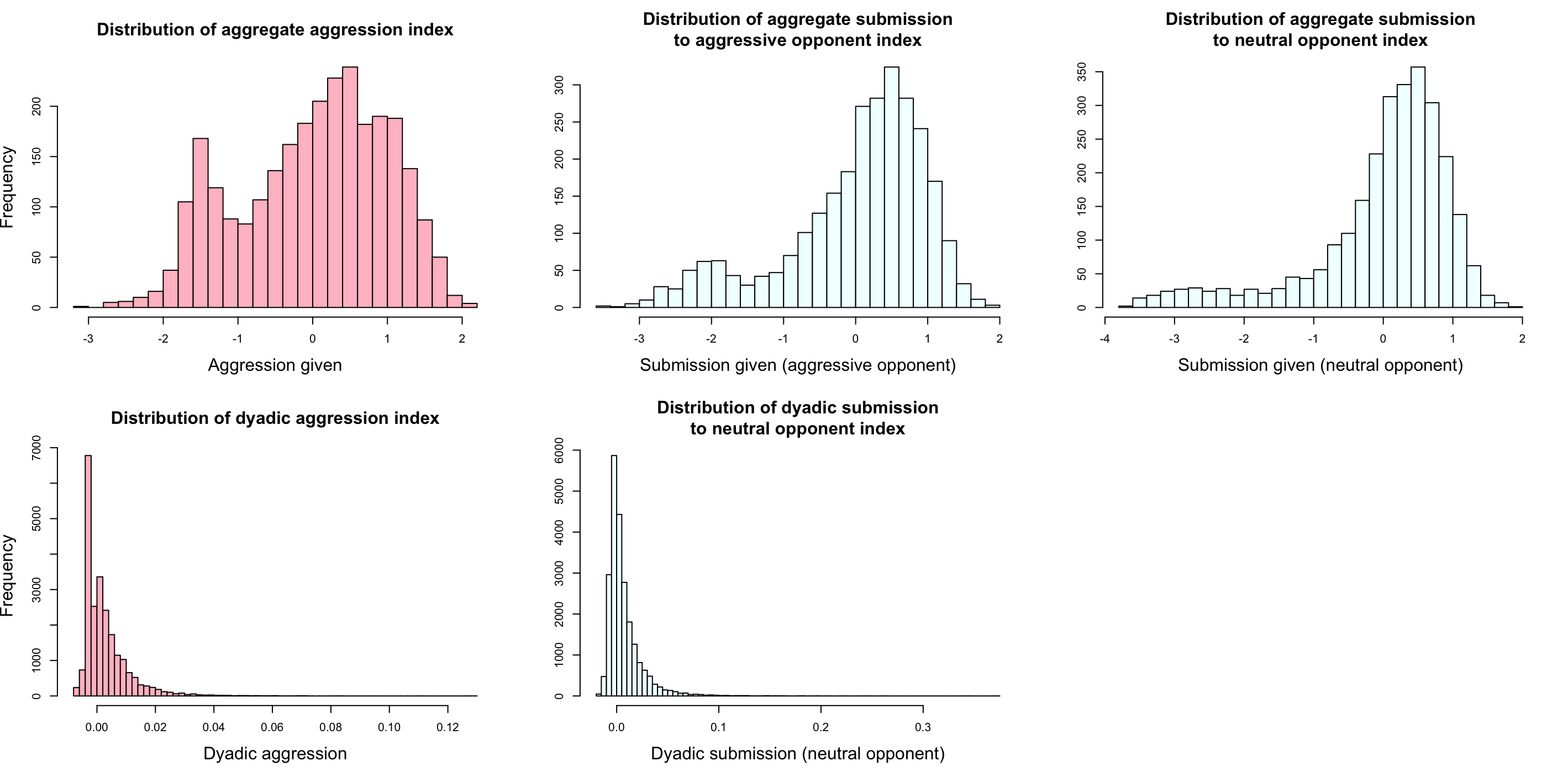

**Supplementary Figure S1**. Distribution of agonistic behavior indices. Distributions in pink are obtained from aggressive-submissive interactions (AS); distributions in blue are obtained from neutral submissive interactions (NS).
